# Large-Scale Neural Network Compensation Underlying Camouflaging in Trait Autism and Its Potential Mental Health Costs

**DOI:** 10.1101/2025.08.22.671791

**Authors:** Han Guo, Xiaobing Chen, Aihua Zhou, Juan Kou, Yi Lei, Keith M. Kendrick, Lei Xu

**Author notes:** Corresponding author: Lei Xu Address: No. 5, Jing’an Road, Jinjiang District, Chengdu, China. 610066.

## Abstract

Social camouflaging refers to strategies to hide or compensate for social difficulties, often at a significant mental health cost, and is particularly prevalent in autism. The large-scale neural network mechanisms underlying this adaptation remain poorly understood. This study aimed to identify these neural underpinnings and their link to other potential mental health issues. Using a dimensional approach, we recruited 110 healthy young adults who completed self-report questionnaires measuring autistic traits and camouflaging as well as depression and anxiety, and underwent resting-state fMRI scans. Using the 300-node Seitzman atlas encompassing 13 functional networks, we examined the interaction between camouflaging and autistic traits on brain network connectivity. Results showed that among individuals with higher autistic traits, greater camouflaging was associated with hyperconnectivities between the Default Mode (DMN) and Cingulo-Opercular Networks (CON), and within the CON. Crucially, DMN-CON hyperconnectivity mediated the relationship between camouflaging and potential mental health cost (i.e., depression and anxiety scores) but only in individuals with higher autistic traits. These findings reveal a specific neuro-compensatory mechanism underlying autistic camouflaging involving self-referential and executive control systems and provide a neurobiological explanation for its potential mental health burden, highlighting the need for societal changes that reduce the pressure for such adaptations.

**Lay Abstract:** Social camouflaging refers to strategies autistic people often use to hide their social differences to better adapt to everyday social contexts. While it may help in the short term, this constant effort can be mentally exhausting and is often linked to severe anxiety and depression. However, little is known about how different brain networks work together to enable social camouflaging and might contribute to a greater risk of anxiety and depression. In this study, autistic traits were measured in 110 young healthy adults in addition to questionnaire-based measures of social camouflaging, anxiety, and depression. Additionally, brain activity was recorded using magnetic resonance imaging while they were resting and not doing any task. We found that, in people with higher autistic traits, social camouflaging was linked to stronger connections between the brain network involved in thinking about oneself (self-reflection; Default Mode Network) and the one in charge of maintaining goal-directed behavior (action control; Cingulo-Opercular Network), as well as stronger connections within the latter brain network. These stronger connections between the self-reflection network and action control network partly explained why camouflaging may be associated with higher levels of anxiety and depression. In short, autistic camouflaging seems to rely on a cognitively demanding neural compensatory process, which can take a toll on mental health. Understanding this underscores the importance of creating a more accepting society to support the well-being of autistic individuals.

## 1 Introduction

Atypical social communication and interaction, as well as restricted, repetitive behaviors or interests, are core symptoms of autism in clinical contexts (DSM-5-TR, 2022). Autistic traits share these qualitative features and are continuously distributed across the general population (Constantino & Todd, 2003; Gökçen et al., 2014), and even non-clinical individuals high in such traits often face similar social challenges (Ingersoll, 2010; Jobe & White, 2007). To better adapt to the demands of everyday predominantly neurotypical social contexts, some autistic individuals or those high in autistic traits consciously or unconsciously adopt strategies—known as social camouflaging—to minimize the visibility of their differences and challenges (Hull et al., 2017; Livingston & Happé, 2017). While camouflaging may facilitate short-term social acceptance, it is also linked to diagnostic challenges (e.g., delayed, missed, or incorrect diagnosis; McQuaid et al., 2021) and adverse mental health outcomes (e.g., anxiety, depression, and suicide ideation; Khudiakova et al., 2024; Ross et al., 2023).

Previous qualitative and quantitative studies have shown that social camouflaging is a multidimensional construct, comprising three core components: masking (hiding autistic traits), compensation (using cognitive strategies to overcome social difficulties), and assimilation (striving to blend in; Hull et al., 2017; Livingston et al., 2019). Based on this framework, the Camouflaging Autistic Traits Questionnaire (CAT-Q; Hull et al., 2019) was developed and is now widely used to measure social camouflaging. Recent theoretical studies indicated that camouflaging is not specific to autism, but reflects a more general human tendency for impression management (Ai et al., 2022; Schneid & Raz, 2020). However, given that the core features of ASD involve social communication difficulties, social camouflaging in autistic individuals still exhibits unique characteristics (e.g., motivations, neurocognition, and consequences; see details in Ai et al., 2022). Despite these insights, the autistic-specific neural mechanisms supporting social camouflaging and their implications for psychological well-being remain poorly understood.

Several empirical studies have demonstrated that the main social symptoms of autism can be partially alleviated through functional compensation, typically by utilizing alternative compensatory neural pathways (Fishman et al., 2018; Pereira et al., 2024; Xu et al., 2022). As a form of social compensation, camouflaging in autism may also involve specific compensatory cognitive–neural mechanisms. At the cognitive level, camouflaging in autism has been linked to enhanced executive functioning (Hull, Petrides, et al., 2021; Lai et al., 2017; Livingston et al., 2019). At the neural level, camouflaging in autistic females has been associated with increased activity in the medial prefrontal cortex (mPFC), a core node of the Default Mode Network (DMN), during self-representation (Lai et al., 2019), as well as with the excitation-inhibition balance within this region during resting-state (Trakoshis et al., 2020). Functional and structural connectivity patterns related to camouflaging have also been observed within circuits implicated in reward, emotion, and memory retrieval (Walsh et al., 2022). These findings suggest that camouflaging in autism is not supported by separate brain regions but by interconnected neural circuits, highlighting the need for further investigation of the compensatory mechanisms involved from a large-scale network perspective.

To address this, the current study analyzed resting-state fMRI data from 110 healthy adults with a dimensional approach using the 300-node Seitzman atlas (Seitzman et al., 2020), which incorporates subcortical and cerebellar regions into 13 functional networks, to examine large-scale brain networks associated with the camouflaging × autistic trait interaction in order to establish potential mechanisms specific to camouflaging in individuals exhibiting higher autistic traits. Based on previous neuroimaging findings, we primarily focused on the DMN and networks supporting executive function (e.g., fronto-parietal network, FPN; cingulo-opercular network, CON; Wallis et al., 2015; Wood & Nee, 2023), as well as their dynamic switching hub—salience network (SN; Menon, 2018; Menon & Uddin, 2010; Uddin, 2015), and their systematic interactions with other large-scale networks.

Social camouflaging in autistic individuals often occurs without full insight into neurotypical social norms and involves distinct ‘high-cost’ psychological mechanisms to compensate for the core symptom (Ai et al., 2022). As a result, autistic individuals frequently experience fatigue and feelings of inauthenticity from camouflaging compared to non-autistic individuals (Hull et al., 2017), which can contribute to worse mental health (Cassidy et al., 2020; Hull, Levy, et al., 2021; Ross et al., 2023; van der Putten et al., 2024). Thus, these specific compensatory mechanisms might operate in a ‘high-cost’ manner and underlie the adverse mental health consequences of camouflaging in autistic individuals, providing a rationale to examine brain network mechanisms as potential mediators.

Taken together, the present study aimed to investigate the specific neural network mechanisms supporting camouflaging in individuals with higher autistic traits and their association with its potential mental health costs. We hypothesized that: enhanced inter- or intra-network connectivity (e.g., within the DMN, FPN, CON, and SN) would facilitate compensatory camouflaging in individuals with higher autistic traits, and these high-cost compensatory pathways would mediate the relationship between camouflaging and adverse mental health outcomes in the context of higher autistic traits.

## 2 Materials and methods

### 2.1 Participants

The final sample for the current study consisted of 110 right-handed healthy adults (76 females; age range = 17-28 years, *M* = 19.93, *SD* = 1.54). These participants were recruited from an initial cohort of 124 college students in China. Fourteen individuals were excluded from the analysis due to excessive head motion (>3 mm translation or >3° rotation) or missing data on the Camouflaging of Autistic Traits Questionnaire (CAT-Q). All participants were free of any self-reported history of diagnosed mental or neurological disorders. The study protocol was approved by the local ethics committee at the Institute of Brain and Psychological Science, Sichuan Normal University (Approval No. 2024LS028) and was conducted in accordance with the latest revision of the Declaration of Helsinki. All participants provided written informed consent prior to the experiment (parental consent was obtained for one participant under the age of 18) and received financial compensation.

### 2.2 Scales

#### 2.2.1 Camouflaging of Autistic Traits Questionnaire (CAT-Q)

The Chinese version of the Camouflaging of Autistic Traits Questionnaire (CAT-Q; Hull et al., 2019) was used to assess camouflaging. The CAT-Q is a 25-item self-report instrument on which participants rate their behaviors on a 7-point scale (from 1 = strongly disagree to 7 = strongly agree). Five items are reverse-scored, and a total score is calculated by summing all item scores, with higher scores indicating greater use of camouflaging. The Cronbach’s alpha for the scale in the current sample was 0.829, demonstrating good internal consistency.

#### 2.2.2 Autism-Spectrum Quotient (AQ)

The Chinese version of the Autism-Spectrum Quotient (AQ; Baron-Cohen et al., 2001) was used to assess autistic traits. The AQ is a 50-item self-report instrument on which participants rate statements on a 4-point scale (from 1 = definitely agree to 4 = definitely disagree). For scoring, a binary system is used: each item is converted to 0 or 1 depending on whether the response reflects the presence of autistic traits. A total score is then calculated by summing all item scores, with higher scores indicating a greater level of autistic traits. The Cronbach’s alpha for the scale in the current sample was 0.715, demonstrating acceptable internal consistency.

#### 2.2.3 Trait Anxiety Inventory (TAI)

The Chinese version of the Trait Anxiety Inventory (TAI) subscale from the State-Trait Anxiety Inventory (STAI; Spielberger et al., 1971) was used to assess trait anxiety. The TAI is a 20-item self-report instrument on which participants rate how they generally feel on a 4-point scale (from 1 = almost never to 4 = almost always). After reverse-scoring the ten appropriate items, a total score is calculated by summing all item scores, with higher scores indicating greater trait anxiety. The Cronbach’s alpha for the scale in the current sample was 0.885, demonstrating excellent internal consistency.

#### 2.2.4 Beck Depression Inventory (BDI)

The Chinese version of the Beck Depression Inventory (BDI-2; Beck et al., 1996) was used to assess depressive symptoms. The BDI-2 is a 21-item self-report instrument on which participants rate the severity of their symptoms on a 4-point scale (from 0 to 3). A total score is calculated by summing the ratings, with higher scores indicating greater levels of depression. The Cronbach’s alpha for the scale in the current sample was 0.910, demonstrating excellent internal consistency.

### 2.3 MRI data acquisition and preprocessing

MRI data were collected on a 3T Prisma MR scanner (Siemens, Erlangen, Germany) with a 64-channel head-neck coil at the Centre for MRI Research, Sichuan Normal University. During the scan, participants were instructed to rest with their eyes open, fixating on a blank screen, and remain as still as possible. Foam padding was used to minimize head motion. Resting-state functional MRI (rs-fMRI) data were acquired using a T2*-weighted echo-planar imaging (EPI) sequence (repetition time = 2000 ms, echo time = 30 ms, flip angle = 90°, FOV = 224 mm × 224 mm, 62 axial slices, slice thickness = 2.0 mm, slice gap = 0.3 mm, voxel size = 2 × 2 × 2.3 mm, multi-band factor = 2). Each run lasted approximately 8 minutes and consisted of 242 volumes. A high-resolution T1-weighted anatomical image was also acquired to improve the normalization of the functional images (repetition time = 2530 ms, echo time = 2.98 ms, flip angle = 7°, matrix size = 224 × 256, 192 sagittal slices, slice thickness = 1.0 mm, voxel size = 0.5 × 0.5 × 1 mm).

Preprocessing of resting-state fMRI data was performed using the Data Processing Assistant for Resting-State fMRI (DPARSF v5.4_230110; Yan & Zang, 2010), a toolbox based on SPM12 and DPABI (Yan et al., 2016). The initial five volumes of each run were discarded to allow for scanner signal stabilization. The remaining functional images underwent slice-timing correction. Structural T1 images were segmented and bias-corrected. Functional images were then co-registered to the individual’s T1 image before being normalized to Montreal Neurological Institute (MNI) standard space and resampled to an isotropic 2 × 2 × 2 mm³ voxel size. Subsequently, the data were cleaned by regressing out several nuisance covariates, including 24 motion parameters derived from realignment, as well as mean signals from white matter, cerebrospinal fluid, and the whole brain (global signal). The residual data were then linearly detrended, band-pass filtered (0.01–0.1 Hz), and spatially smoothed with a 6-mm FWHM Gaussian kernel.

### 2.4 Functional Network Analysis

#### 2.4.1 Functional Network Construction

Functional brain networks were constructed for each participant by first defining connectivity at the level of regions of interest (ROIs) and subsequently summarizing these into network-level metrics. To establish ROI-level connectivity, each participant’s brain was parcellated into 300 ROIs using the Seitzman et al. (2020) atlas. This atlas assigns ROIs to one of 13 functional networks (e.g., DMN, FPN, CON, SN; see Supplementary Table S1 for a full list and MNI coordinates), excluding the ‘unassigned’ category. After extracting the mean time series from each ROI, a 300 × 300 connectivity matrix was generated for each participant by computing Fisher’s Z-transformed Pearson’s correlations between all ROI pairs. To focus our analysis on synchronous neural activity and avoid the ambiguous interpretation of negative correlations, all negative correlation values were set to zero. To ensure the robustness of our findings, this process was repeated across multiple sparsity thresholds (10%, 30%, 50%, 70%, and 90%), retaining only the strongest positive connections at each level.

These detailed ROI-level matrices were then summarized to create the network-level metrics used in our primary analysis. Specifically, intra-network connectivity was calculated by averaging the correlation values among all unique node pairs within a given network, while inter-network connectivity was derived from the average correlations of all node pairs connecting two different networks. This procedure yielded a 13 × 13 summary matrix of mean functional connectivity for each participant at each sparsity threshold. The diagonal elements of this matrix represent intra-network connectivity, and the off-diagonal elements represent inter-network connectivity.

#### 2.4.2 Network-based Statistical Analysis

Our primary statistical analysis focused on the network-level connectivity matrices. A General Linear Model (GLM) was conducted to test for the effects of camouflaging (CAT-Q), autistic traits (AQ), and their interaction on each of the intra- and inter-network connections, while controlling for age and sex. The resulting p-values for the 13 intra-network and 78 inter-network connections were independently corrected for multiple comparisons using the False Discovery Rate (FDR) procedure (*p*FDR < 0.05). For robustness, we prioritized network connections that remained significant across all tested sparsity thresholds.

To visualize and localize the specific connections and flexible hubs driving any significant network-level effects, we subsequently utilized the Network-Based Statistics (NBS) toolbox (Zalesky et al., 2010). For any significant network-level finding, NBS was used to identify the specific subnetwork of edges significantly associated with our predictors (pFWE < 0.05, corrected at the component level after 10,000 permutations). All statistical analyses were performed in MATLAB R2018a (The MathWorks, Inc.).

### 2.5 Associations between Functional Network and Mental Health Outcomes

To test whether the identified neural pathways mediated the relationship between social camouflaging and mental health, we conducted moderated mediation analyses. Before that, partial correlation analyses were performed to first identify which of the robustly identified neural markers (i.e., those showing a significant interaction effect across all sparsity thresholds) were also significantly associated with our mental health measures (depression and anxiety). These analyses controlled for age and sex, and p-values were corrected for multiple comparisons using FDR. Based on these results, any neural pathway significantly associated with mental health was then tested in a moderated mediation model using the PROCESS macro (Hayes, 2022) for SPSS (Model 7). In these models, we tested the indirect effect of camouflaging (CAT-Q; independent variable, X) on mental health outcomes (anxiety or depression; dependent variable, Y) via the significant neural pathway (mediator, M). This indirect pathway was specified to be moderated by autistic traits (AQ; moderator, W). The analysis was based on 5,000 bootstrap samples, and 95% confidence intervals (CIs) were generated to test the significance of the conditional indirect effects and the index of moderated mediation. To further probe the significant moderated mediation, the Johnson-Neyman (J-N) technique was employed to identify the precise range of autistic traits (the moderator) for which the conditional effect of camouflaging on the neural mediator was statistically significant. The resulting J-N plot was generated using the Jnplots package (Toyama, 2024) in the R statistical environment (version 4.4.1).

## 3. Results

### 3.1 Behavioral Results: Camouflaging is Associated with Autistic Traits and Poorer Mental Health

Descriptive statistics and correlations for all study variables are presented in Table 1. Pearson correlation analysis revealed a significant positive association between camouflaging (CAT-Q) and autistic traits (AQ; *r* = 0.286, *p =* 0.002), indicating that individuals with higher autistic traits reported more camouflaging behaviors. Camouflaging was also significantly associated with poorer mental health, showing positive correlations with both depression (BDI; *r* = 0.246, *p =* 0.010) and trait anxiety (TAI; *r* = 0.310, *p =* 0.001). Furthermore, autistic traits (AQ) were also significantly correlated with depression and anxiety levels (see Table 1). These behavioral results confirm the core associations between social camouflaging, autistic traits, and mental health in our sample, providing a foundation for investigating their underlying neural mechanisms.

**Table 1:** Descriptive statistics and correlations for the study variables.

| Variables | Mean ± SD | 1 | 2 | 3 | 4 | 5 |
| --- | --- | --- | --- | --- | --- | --- |
| 1. Gender | 76 F / 34 M |  |  |  |  |  |
| 2. Age | 19.93 ± 1.54 | -0.302** | | | | |
| 3. CAT-Q | 108.55 ± 15.31 | 0.271** | -0.147 | | | |
| 4. AQ | 24.17 ± 6.02 | 0.216* | -0.17 | 0.286** | | |
| 5. TAI | 46.45 ± 9.06 | 0.241* | -0.119 | 0.31** | 0.584** | |
| 6. BDI | 11.50 ± 8.80 | 0.166 | -0.14 | 0.246** | 0.383** | 0.679** |

\* $p < 0.05$ , \*\* $p < 0.01$ for Pearson correlation. CAT-Q, Camouflaging of Autistic Traits Questionnaire; AQ, Autism-Spectrum Quotient; TAI, Trait Anxiety Inventory; BDI, Beck Depression Inventory; M, male; F, female; SD, standard deviation.

### 3.2 Brain Network Connectivity Results

#### 3.2.1 Selection of Sparsity Thresholds for Network Construction

To construct the functional connectivity matrices, all negative connections were excluded, and multiple sparsity thresholds (10%, 30%, 50%, 70%, and 90%) were applied to the positive connectivity matrices to ensure the robustness of our results. The proportion of zero-value entries in each participant’s network-level connectivity matrix at each threshold was evaluated. A network-level connection (e.g., DMN-CON inter-network FC) was defined as zero if all constituent ROI-to-ROI correlations were set to zero by the thresholding procedure. As shown in Supplementary Table S2, the 10% sparsity threshold resulted in an excessive proportion of zero-value entries, indicating a significant loss of network information and potential fragmentation of the network topology. Consequently, this threshold was excluded from further statistical analyses. The remaining sparsity thresholds (30%, 50%, 70%, and 90%) yielded an acceptable proportion of zero-value entries and were retained for all subsequent analyses. To maximize the signal-to-noise ratio and clearly delineate the most critical neural circuits, we primarily report the results from the 30% sparsity threshold in the main text, and the rest of the sparsity results are presented in the supplementary material.

#### 3.2.2 Interaction Effect of Camouflaging and Autistic Traits on Network Connectivity

A General Linear Model (GLM) was used to examine the effects of social camouflaging (CAT-Q), autistic traits (AQ), and their interaction on the 13 intra-network and 78 inter-network functional connections, while controlling for age and sex. The analysis revealed a significant and robust interaction between CAT-Q and AQ scores (see Figure 1A & B). Across all tested sparsity thresholds (30%, 50%, 70%, and 90%), the interaction term significantly and positively predicted functional connectivity between the Default Mode Network (DMN) and the Cingulo-Opercular Network (CON) (30% sparsity: *B* = 0.010, *t* = 3.401, *p*FDR = 0.037; 50% sparsity: *B* = 0.008, *t* = 3.341, *p*FDR = 0.023; 70% sparsity: *B* = 0.007, *t* = 3.386, *p*FDR = 0.026; 90% sparsity: *B* = 0.005, *t* = 3.336, *p*FDR = 0.046). The interaction also consistently predicted intra-network connectivity within the CON (30% sparsity: *B* = 0.014, *t* = 3.573, *p*FDR = 0.007; 50% sparsity: *B* = 0.011, *t* = 3.320, *p*FDR = 0.008; 70% sparsity: *B* = 0.010, *t* = 3.348, *p*FDR = 0.015; 90% sparsity: *B* = 0.009, *t* = 2.920, *p*FDR = 0.028). Simple slopes analyses revealed that among individuals with higher autistic traits, greater social camouflaging was associated with stronger DMN-CON inter-network and CON intra-network functional connectivity (see Figure 2C for details).

**Figure 1:**
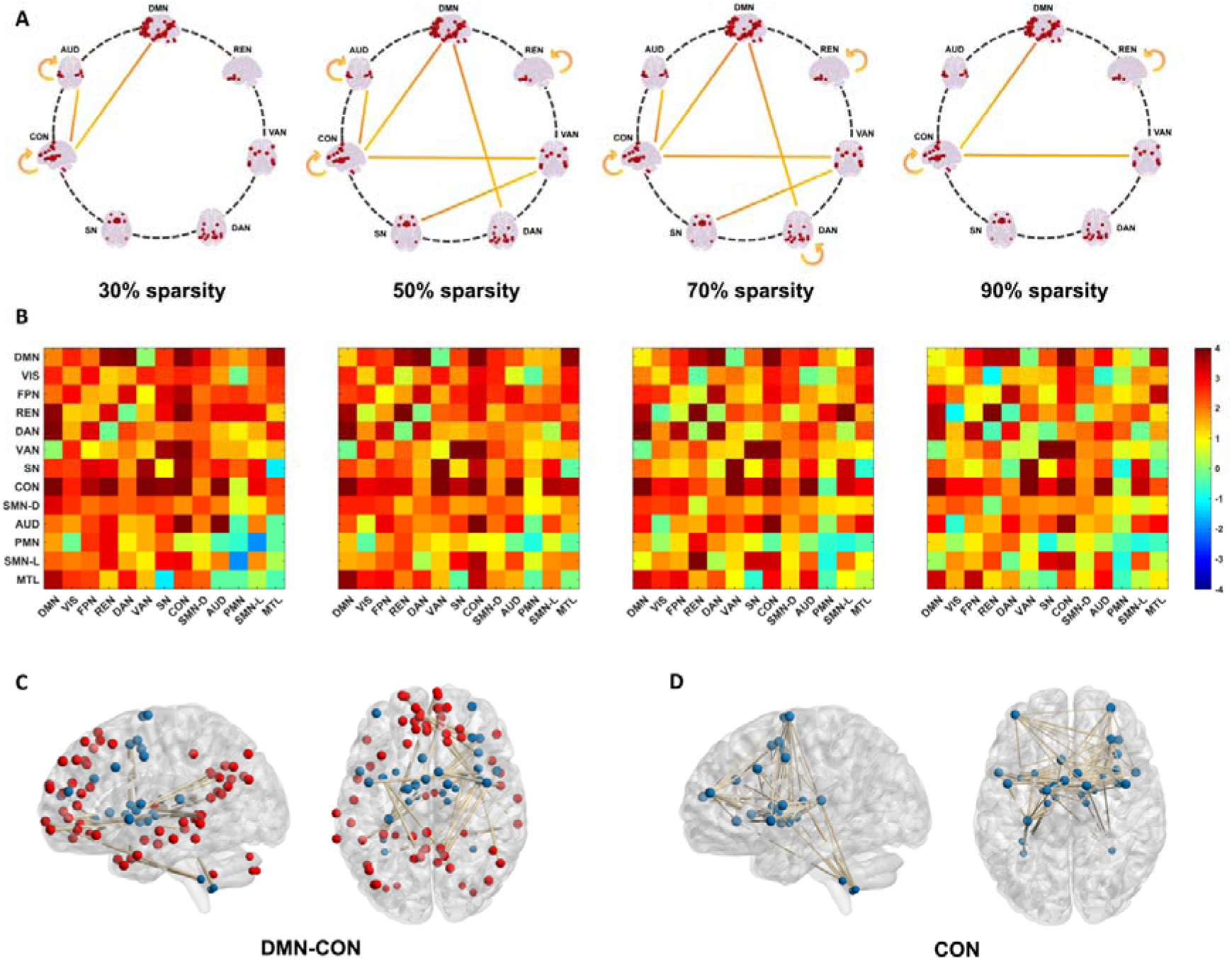
Neural Correlates of the Camouflaging × Autistic Traits Interaction. (A) Functional connections showing a significant interaction effect (*p*FDR < 0.05) across four sparsity thresholds. Robust effects were found for DMN-CON between-network and CON intra-network connectivity. (B) T-value matrices of the interaction effect at each sparsity threshold. (C-D) Sub-networks identified by Network-Based Statistics (*p* < 0.05, FWE-corrected), showing enhanced connectivity between (C) the Default Mode Network (DMN; red nodes) and Cingulo-Opercular Network (CON; blue nodes), and (D) within the CON. Brain nodes are displayed in sagittal (left) and axial (right) views.

**Figure 2:**
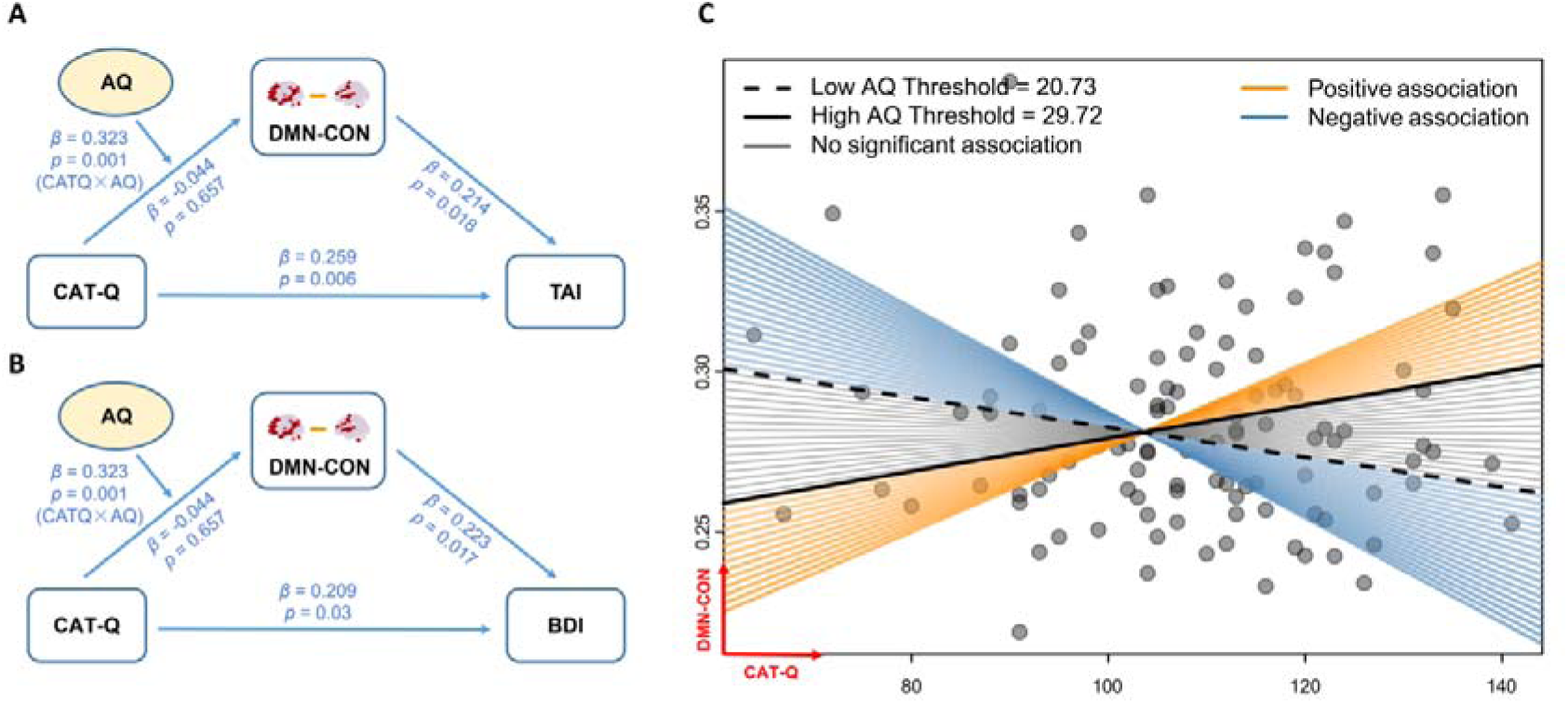
(A-B) Moderated mediation models showing the indirect effect of social camouflaging (CAT-Q) on (A) trait anxiety (TAI) and (B) depression (BDI) through DMN-CON functional connectivity, moderated by autistic traits (AQ). Unstandardized regression coefficients are shown for each path. (C) Johnson-Neyman (J-N) plot illustrating the conditional effect of social camouflaging on DMN-CON functional connectivity across the range of autistic trait scores. The orange and blue regions represent significant positive and negative associations, respectively, while the gray region indicates non-significance. The solid and dashed vertical lines indicate the AQ values at which the effect becomes or ceases to be statistically significant.

In addition to this primary finding, other significant effects were observed at specific sparsity thresholds (see Supplementary Table S3). At the 50% and 70% thresholds, the interaction term also significantly predicted connectivity between the DMN and the Dorsal Attention Network (DAN), between the Ventral Attention Network (VAN) and both the Cingulo-opercular Network (CON) and Salience Network (SN), as well as intra-network connectivity within the Reward Network. Regarding the main effect of CAT-Q, it was found to negatively predict intra-network connectivity within the REN at the 70% sparsity threshold (*B* = -0.011, *t* = -2.973, *p*FDR = 0.024) and robustly predicted lower intra-network connectivity within the Auditory Network (AUD) across all thresholds (*B*s ranged from -0.017 to -0.018; *t*s ranged from -3.062 to -3.758; all *p*FDR < 0.036). No significant main effects of AQ scores were found.

#### 3.2.3 The Specific Sub-network for Compensation

To pinpoint the specific neuroanatomical connections driving the robust network-level findings, we employed Network-Based Statistics (NBS) and focused on the 30% sparsity threshold to isolate the strongest associations and clearly reveal the key neural circuits.

The NBS analysis on DMN-CON between-network connections identified a single significant sub-network positively associated with the CAT-Q × AQ interaction (*p*FWE < 0.05). This compensatory circuit consisted of 28 edges connecting 28 nodes. It was characterized by enhanced connectivity between core hubs of the DMN—including the pregenual Anterior Cingulate Cortex, Precuneus, Posterior Cingulate Cortex, and medial/inferior Orbitofrontal Cortex—and key regions of the CON, such as the Insula, Putamen, Thalamus, and Supplementary Motor Area (Figure 1C). Similarly, the NBS analysis on CON intra-network connections revealed a single, densely connected sub-network also significantly and positively associated with the CAT-Q × AQ interaction (*p*FWE < 0.05). This sub-network was extensive, encompassing all nodes within the Cingulo-Opercular Network (144 edges and 30 nodes), suggesting that a global increase in intrinsic CON coherence serves as a key compensatory mechanism (Figure 1D).

### 3.3 The Mediating Role of Compensatory Neural Pathways in Mental Health

To identify a candidate neural mediator, we first examined the correlations between our robustly identified neural markers (DMN-CON inter-network FC and CON intra-network FC at 30% sparsity) and mental health measures. After controlling for age and sex, partial correlation analyses revealed that DMN-CON connectivity was significantly and positively correlated with both trait anxiety (*r =* 0.222, *p*FDR = 0.042) and depression (*r =* 0.228, *p*FDR = 0.042). In contrast, the relationship between CON intra-network connectivity and mental health measures was non-significant (*r*s < 0.121; *p*FDR > 0.284).

Therefore, DMN-CON connectivity was carried forward as the mediator in our moderated mediation models. The analysis revealed a significant overall index of moderated mediation for both the anxiety model (Index = 0.069, 95% CI = [0.011, 0.145]; see Figure 2A) and the depression model (Index = 0.072, 95% CI = [0.014, 0.155]; see Figure 2B). This indicates that the indirect effect of camouflaging on mental health via DMN-CON connectivity was conditional upon an individual’s level of autistic traits. The Johnson-Neyman analysis further specified the boundaries of this conditional effect. It revealed that for individuals with AQ scores above 29.72 (the top 15.46%), camouflaging was significantly and positively associated with DMN-CON connectivity (*B* = 0.254, *p* < 0.05). Conversely, for individuals with AQ scores below 20.73 (the bottom 28.12%), camouflaging was significantly and negatively associated with DMN-CON connectivity (see Figure 2C). After accounting for this mediated pathway, the direct effects of camouflaging on both anxiety (*B* = 0.259, *t* = 2.801, *p* = 0.006) and depression (*B* = 0.209, *t* = 2.205, *p* = 0.030) remained significant. Detailed regression results are presented in Supplementary Table S4 (see Supplementary Tables S5-S7 for results at other sparsity thresholds).

## 4. Discussion

This study aimed to elucidate the large-scale brain networks underlying social camouflaging and its associated mental health costs in individuals with higher autistic traits. The results revealed a specific neural signature of social camouflaging in autistic traits, characterized by enhanced functional connectivity between the Default Mode Network (DMN) and the Cingulo-Opercular Network (CON), as well as increased intra-network connectivity within the CON. Crucially, DMN-CON hyperconnectivity mediated the relationship between camouflaging and adverse mental health outcomes (anxiety and depression) in individuals high in autistic traits, whereas in those with lower autistic traits, camouflaging was associated with DMN–CON hypoconnectivity and correspondingly lower anxiety and depression. These findings reveal a novel network-level ‘high-cost’ compensation mechanism potentially underlying camouflaging in both preclinical and clinical autism, establishing a direct neurobiological link between the compensatory effort involved in camouflaging and its mental health consequences.

### 4.1 Brain Network Compensation of Social Camouflage Associated with Autistic Traits

Our primary and most robust finding was the significant interaction between social camouflaging and autistic traits on both DMN-CON and intra-CON connectivities. The CON supports stable task maintenance, cognitive and physiological readiness for goal-directed behavior (Dosenbach et al., 2008, 2025; Newbold et al., 2021; Zanto & Gazzaley, 2013). Camouflaging-related intra-CON hyperconnectivity in individuals with higher autistic traits might reflect a persistent recruitment of executive control to sustain deliberate evaluation and regulation of their social behaviors in order to align with perceived social expectations. The DMN, implicated in self-referential processing, autobiographical memory retrieval, and prospection (Katsumi et al., 2024; Spreng et al., 2009), provides a foundation for social cognition, including mentalizing about others’ states and establishing shared meaning (Menon,

2023; Schilbach et al., 2008). Typically, these two networks are segregated or anti-correlated to enable flexible switching between internal mentation (including social cognition) and external tasks (Dosenbach et al., 2025; Menon, 2011, 2023). Our finding— that greater camouflaging was associated with stronger coupling between the DMN and CON in individuals high in autistic traits — suggested engagement of typically segregated self-referential and executive control systems, highlighting an increased reliance on executive control to regulate or suppress self-referential processing.

Furthermore, our NBS analysis provided fine-grained support for this interpretation, identifying atypical connections between core social-cognitive hubs of the DMN—such as the mPFC and posterior cingulate cortex (Esménio et al., 2019)—and the CON. This suggests a specific compensatory recruitment of the DMN’s social-referential nodes to achieve the goals of camouflaging. Together, both DMN-CON and intra-CON hyperconnectivity indicated that social camouflaging in autistic individuals may involve more effortful and cognitively costly, goal-directed self-related processing compared to non-autistic individuals. Our findings provided strong empirical neurobiological support for the computational model of camouflaging, which theoretically posits compensatory neurocognitive routes underlying autistic camouflaging (Ai et al., 2022), and further confirmed at the neural network level that self-referential processing (Lai et al., 2019; Trakoshis et al., 2020) and executive functions (Hull, Petrides, et al., 2021; Lai et al., 2017; Livingston et al., 2019), as well as their interaction, are fundamental in supporting these compensatory mechanisms.

Our findings also offered an alternative, more adaptive perspective on the atypical connectivities between DMN and task control networks frequently reported in autism research (Blume et al., 2024; de Lacy et al., 2017; Ilioska et al., 2023; Liu et al., 2025). Rather than considering them as markers of structural or functional impairment (e.g., failed network segregation or cognitive rigidity; Abbott et al., 2016; Menon, 2018; Yang et al., 2023), our findings directly linked these patterns to the strategic behavior of social camouflaging and framed them as evidence of neurally effortful, plastic compensation. Specifically, we propose that autistic camouflaging may reconstrue natural social interaction as a goal-directed task requiring deliberate execution and maintenance, plausibly recruiting task-control CON to sustain top-down modulation of self-referential DMN, thereby aligning the self with the neurotypical social context. Similar results or interpretations have also been reported in previous studies, which indirectly support this perspective, although they did not directly measure camouflaging. For example, enhanced connectivity between DMN and Central Executive Network (CEN, also referred to as FPN) is associated with better metacognitive and social skills in autistic individuals (Blume et al., 2024), where metacognitive strengths may compensate for specific social skill difficulties (Blume et al., 2024; Freeman et al., 2017; Torske et al., 2018); autistic females, who are commonly considered to engage more extensively in camouflaging (Hull et al., 2020) and exhibit stronger DMN–CEN connectivity than both neurotypical females and autistic males (Lawrence et al., 2020).

It should be noted that our findings highlight a specific coupling between DMN and CON rather than FPN/CEN. Whereas the FPN/CEN primarily supports flexible, phasic problem-solving (Chuikova et al., 2025; Cole et al., 2013; Vallesi et al., 2022), the CON supports the stable, sustained maintenance of goal-directed behavior (Dosenbach et al., 2008, 2025; Sadaghiani & D’Esposito, 2015). According to the computational model of camouflaging (Ai et al., 2022), autistic individuals tend to adopt heuristic, low-resolution strategies (e.g., maintaining a fixed smile, imitating actions, or using scripted lines), explaining why autistic camouflaging relies more on ‘action maintenance’ than ‘flexible reasoning’. This distinction is critical as it reveals the specific neurocognitive demands of autistic camouflaging. Our robust finding of enhanced intra-CON connectivity further consolidates this model. In the past, however, the role of the CON in autistic compensation may have been underestimated, possibly due to its ambiguous functional definition and frequent conflation with other networks, such as the salience network. With recent clarifications (Dosenbach et al., 2025), future research is now better positioned to investigate the distinct contributions of the CON.

### 4.2 The Mental Health Cost of Neural Compensation

Another key finding of our study emerged from the moderated mediation analysis, showing that the indirect effect of camouflaging on mental health outcomes, via DMN-CON hyperconnectivity, was significant only in individuals with higher autistic traits. It suggested that although DMN-CON hyperconnectivity may support camouflaging as an adaptive strategy, the chronic engagement of such a neurally effortful process could contribute directly to psychological burden, providing a powerful mechanistic model for why camouflaging is so frequently associated with anxiety and depression in the autistic community (Cassidy et al., 2020; Ross et al., 2023).

A previous study showed that camouflaging in autism is linked to poor mental health through mechanisms such as heightened self-monitoring, cognitive exhaustion, and feelings of inauthenticity (Field et al., 2024). Our findings provide a neural explanation for these negative experiences. The enhanced DMN-CON coupling, involving a network that supports sustained vigilance (CON; Dosenbach et al., 2025; Sadaghiani & D’Esposito, 2015), likely reflects the continuous self-monitoring and high-cost cognitive strategies central to this autistic adaptive strategy. It is this sustained, neurally expensive compensatory activity that plausibly manifests as anxiety and depression. To our knowledge, this is the first study to offer direct neural evidence for this link. It is important to note, however, that this pathway only partially mediated the relationship, suggesting that other factors specific to the experience of camouflaging in autism, such as inauthenticity or internalized stigma (van der Putten et al., 2024), also play crucial roles and warrant future neuro-level investigation.

Intriguingly, the Johnson-Neyman analysis revealed that the neuro-cognitive profile of camouflaging fundamentally differs at lower levels of autistic traits. For these individuals, camouflaging was associated with an opposite neural pattern: DMN-CON hypoconnectivity. This clearer segregation between the DMN and CON may reflect a more efficient, automated form of impression management, where self-referential thought is effectively decoupled from the execution of goal-directed social scripts. This ‘low-cost’ camouflaging may allow individuals to meet social expectations more effortlessly, thereby serving as a protective factor against anxiety and depression. This aligns with findings that associate camouflaging with reduced depression in the general population (Lorenz & Hull, 2024). This stark contrast underscores that camouflaging is not a unitary phenomenon; its underlying neurocognitive processes and associated costs appear to be fundamentally different in the context of higher autistic traits, highlighting the unique and profound burden it represents.

Notably, a recent cross-sectional study suggested that camouflaging in autistic individuals may have some beneficial effects on psychological well-being, but these effects were small (van der Putten et al., 2025). Therefore, autistic camouflaging is more aptly viewed as a double-edged sword. While our cross-sectional design precludes causal claims, our findings provide a compelling hypothesis by elucidating the neural pathway linking camouflaging to its adverse potential mental health consequences, without identifying a neural basis for potential benefits—an issue for future research to address.

### 4.3 Strengths, Limitations, and Future Directions

This study has several strengths, including its robust network construction across multiple sparsity thresholds and the use of a sophisticated moderated mediation model to link brain, behavior, and mental health. However, some limitations must be acknowledged. First, the current sample consists of non-diagnosed individuals, warranting caution when generalizing these findings to individuals with a clinical Autism Spectrum Disorder diagnosis. Second, self-report measures should be complemented by behavioral and clinical assessments in future work. Third, the cross-sectional design, while effective for identifying associations, cannot establish causality; longitudinal studies are needed to unravel the causal pathways linking camouflaging, brain network organization, and mental health. Finally, while the resting-state data reveal an intrinsic network architecture, task-based fMRI is required to test whether this DMN-CON hyperconnectivity is specifically engaged during active social camouflaging.

## 5 Conclusion

This study provides compelling large-scale network evidence for a specific neuro-compensatory mechanism underlying social camouflaging in individuals with higher autistic traits: hyperconnectivities between the default mode and cingulo-opercular networks, as well as intra-cingulo-opercular network. By demonstrating how this neural effort is linked to the well-documented mental health burden of camouflaging, our work offers a powerful neurobiological model for this phenomenon. These findings deepen our understanding of the immense effort expended by autistic individuals to navigate a neurotypical world and underscore the urgent need to drive societal change to reduce the pressure for such costly adaptations, thereby promoting the well-being of all individuals.

## Acknowledgments section

The authors are grateful to all the participants for their enthusiastic involvement in this study. The authors would also like to thank Dr. Meng-Chuan Lai for sharing the Chinese version of the Camouflaging Autistic Traits Questionnaire.

## Author contributions

**Han Guo:** Conceptualization, Methodology, Formal Analysis, Visualization, Writing – Original Draft Preparation, Writing – Review & Editing. **Xiaobing Chen:** Investigation, Data curation, Writing – Review & Editing.**Aihua Zhou:** Investigation, Writing – Review & Editing**. Juan Kou:** Conceptualization, Validation, Writing – Review & Editing. **Yi Lei:** Investigation, Resources, Writing – Review & Editing. **Keith M. Kendrick:** Conceptualization, Validation, Writing – Review & Editing. **Lei Xu:** Conceptualization, Methodology, Supervision, Funding Acquisition, Project Administration, Writing – Original Draft Preparation, Writing – Review & Editing.

## Declaration of conflicting interest

The authors declare that the research was conducted in the absence of any commercial or financial relationships that could be construed as a potential conflict of interest.

## Funding statement

This work was supported by the National Natural Science Foundation of China (grant number NSFC32100893; Sichuan Province Key Research and Development Project (grant number 2023YFWZ0003).

## Ethical approval and informed consent statements

The study protocol was approved by the local ethics committee at the Institute of Brain and Psychological Science, Sichuan Normal University (Approval No. 2024LS028) and was conducted in accordance with the latest revision of the Declaration of Helsinki. All participants provided written informed consent prior to the experiment (parental consent was obtained for one participant under the age of 18) and received compensation.

## Data availability statement

The anonymized dataset is available on reasonable request from the corresponding author.

## Supplementary Information

The online version contains supplementary material available at

## Abbreviations

CAT-Q: Camouflaging Autistic Traits Questionnaire
AQ: Autism-Spectrum Quotient
TAI: Trait Anxiety Inventory
BDI: Beck Depression Inventory
DMN: Default Mode Network
VIS: Visual Network
PFN: Fronto Parietal Network
REN: Reward Network
DAN: Dorsal Attention Network
VAN: Ventral Attention Network
SN: Salience Network
CON: Cingulo-Opercular Network
SMN-D: Dorsal Somatomotor Network
SMN-L: Lateral Somatomotor Network
AUD: Auditory Network
PMN: Parieto-Medial Network
MTL: Medial Temporal Lobe Network.

## Supplementary Material

**Table S1:**
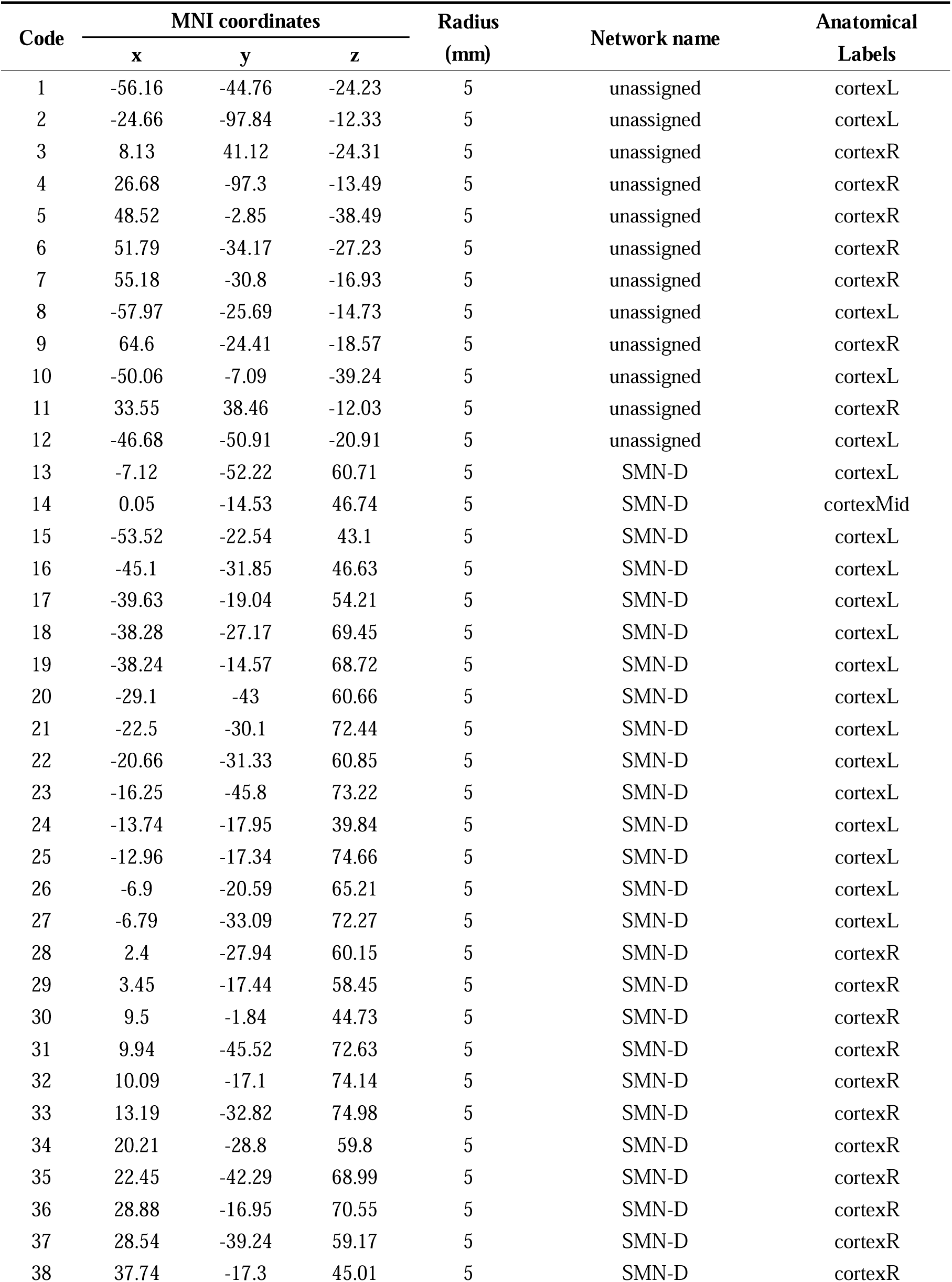
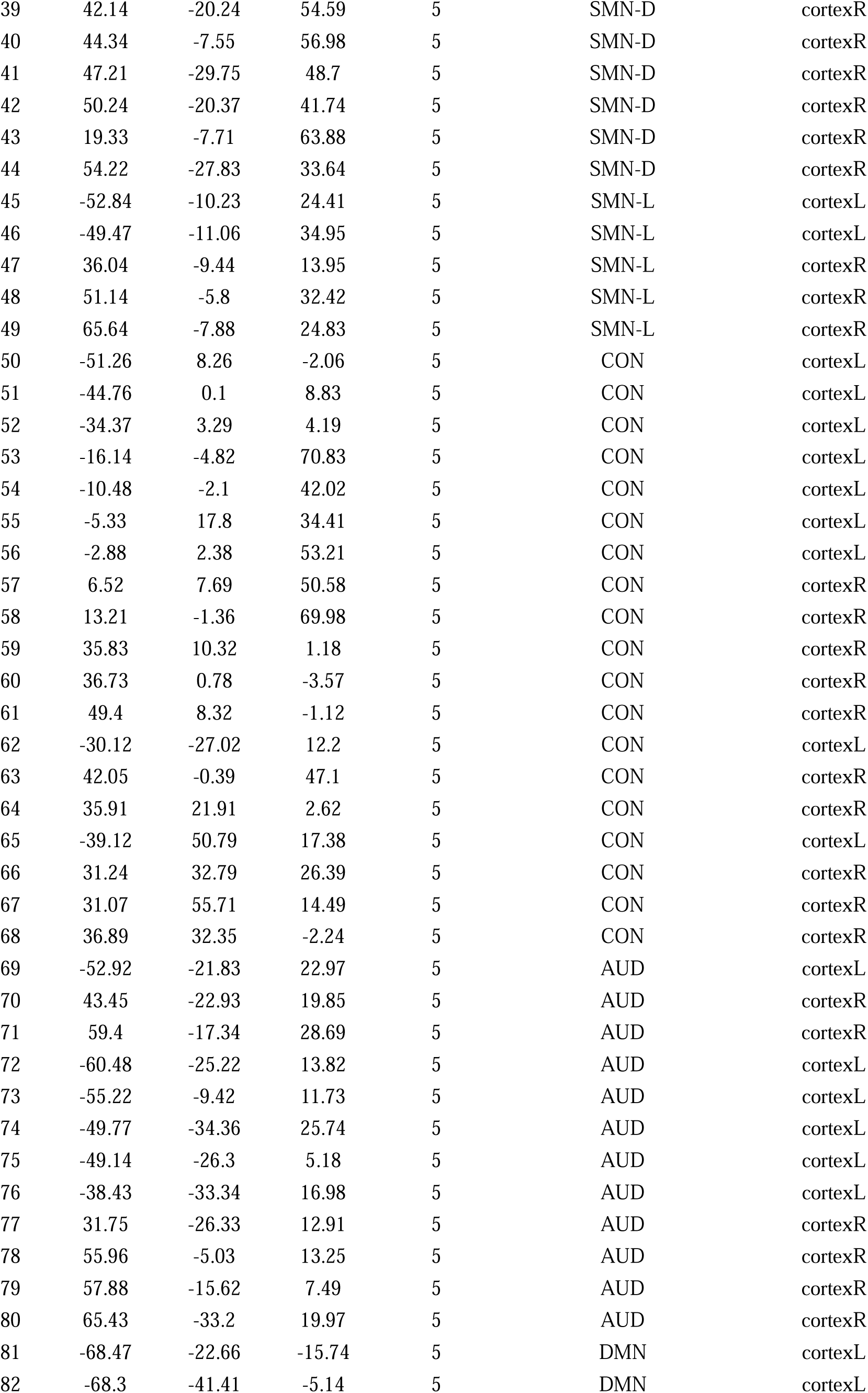
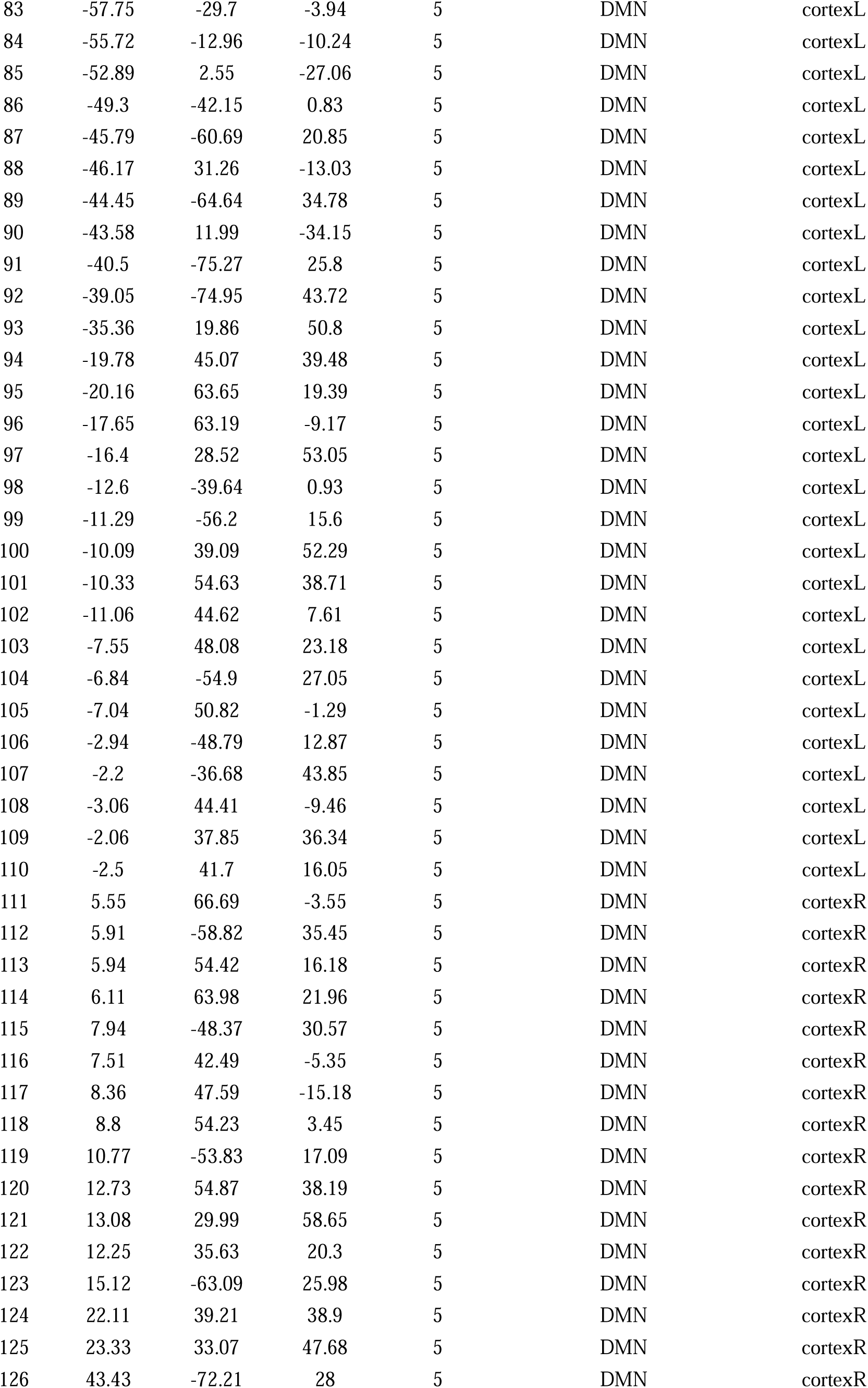
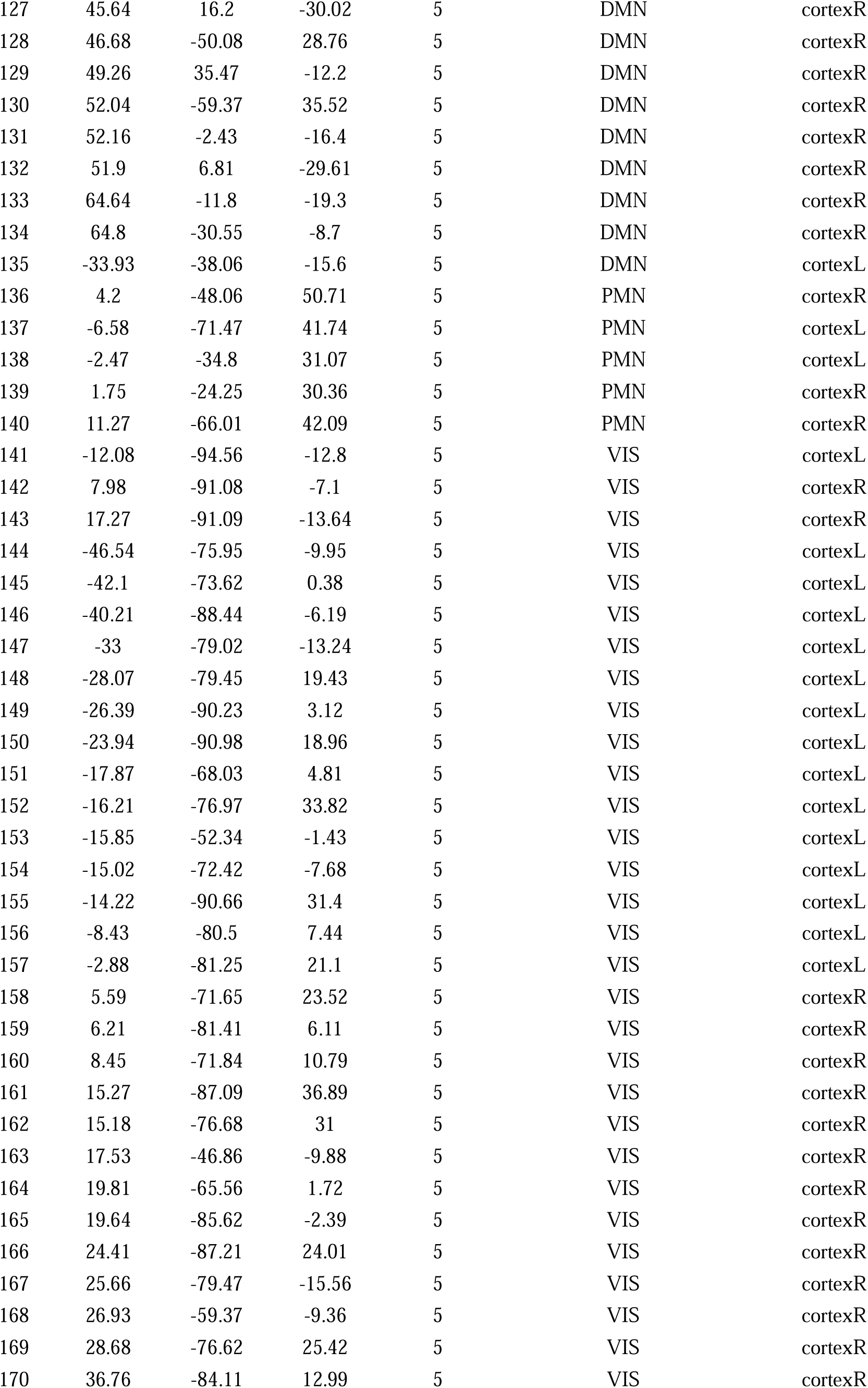
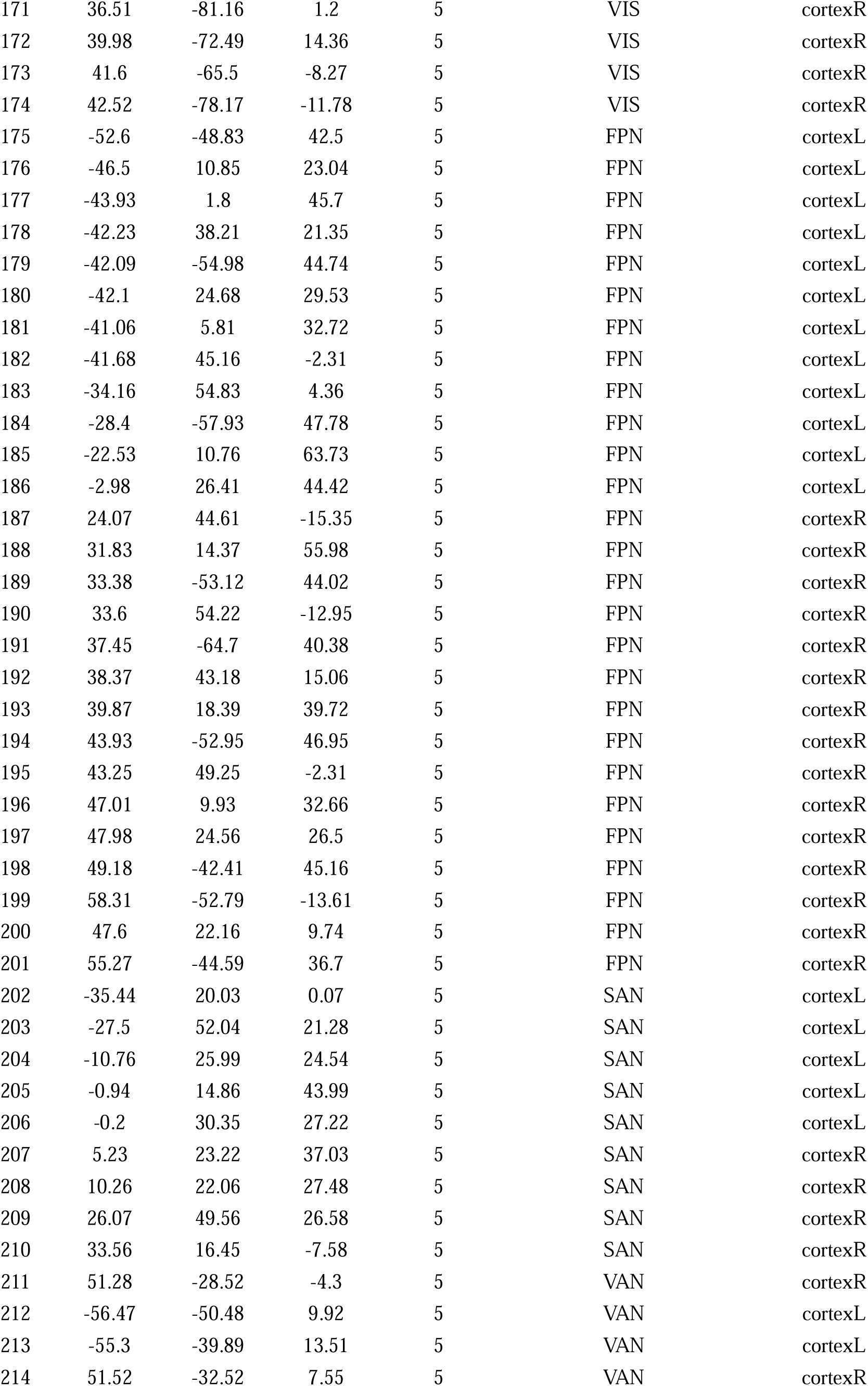
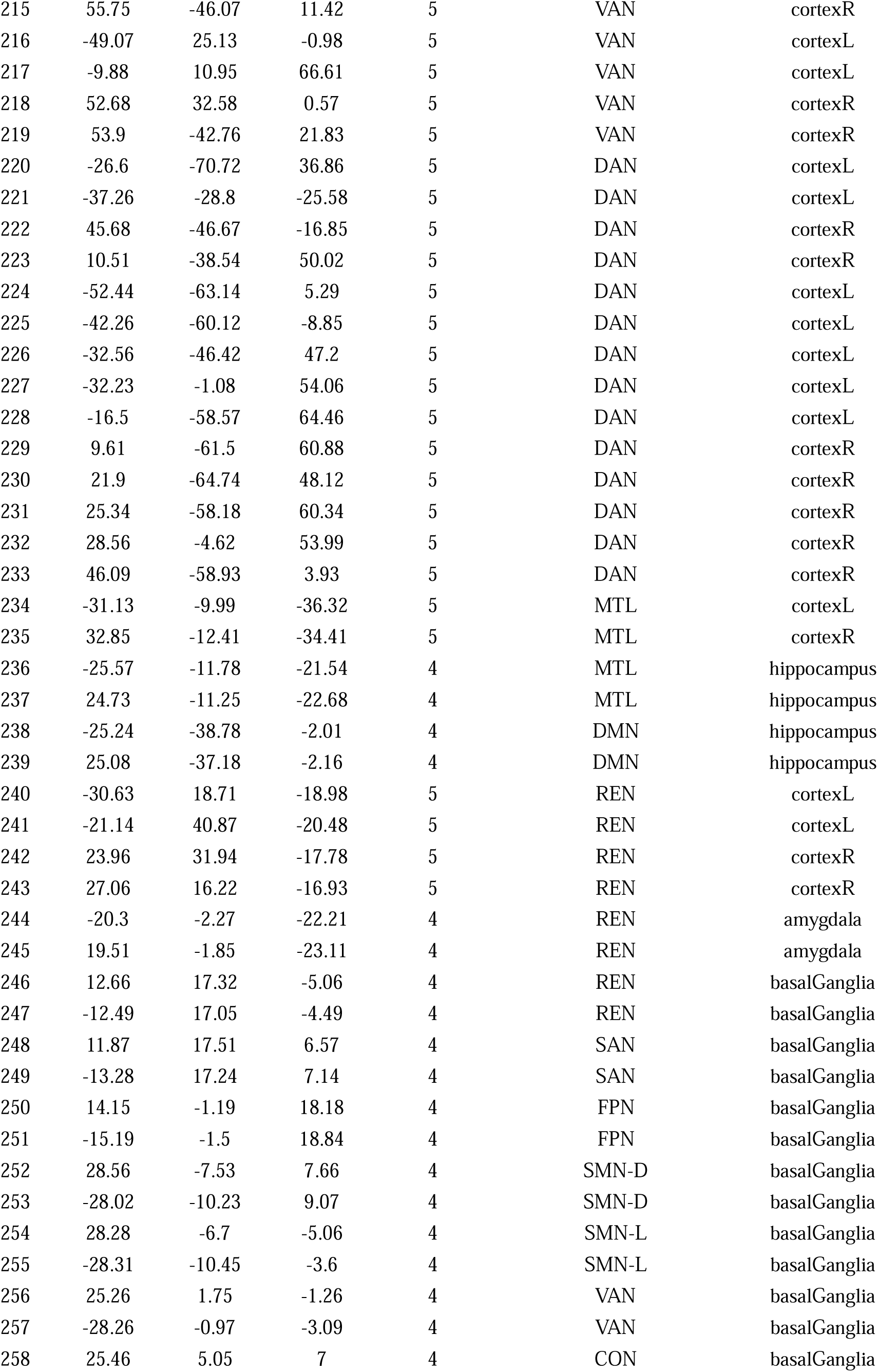
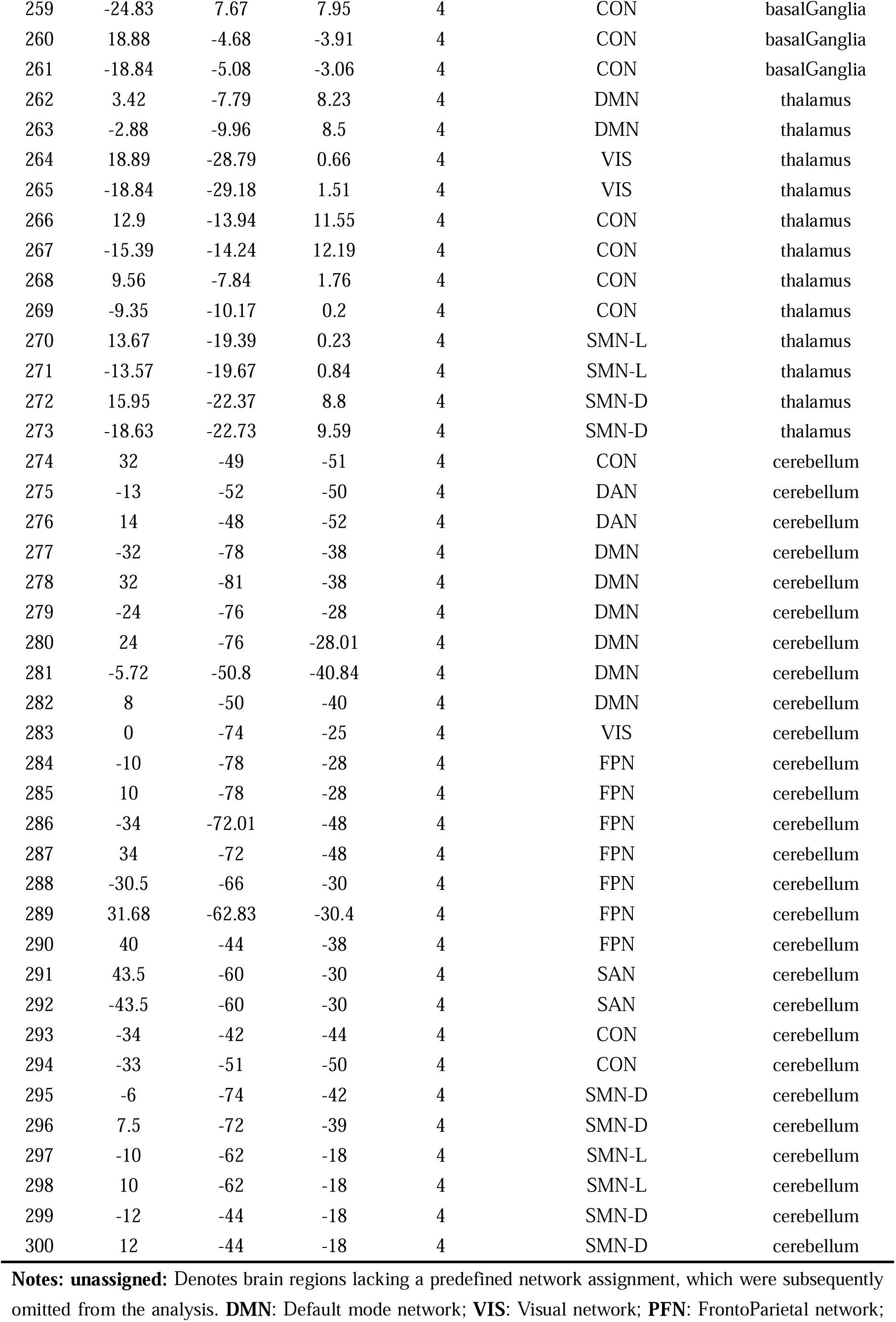
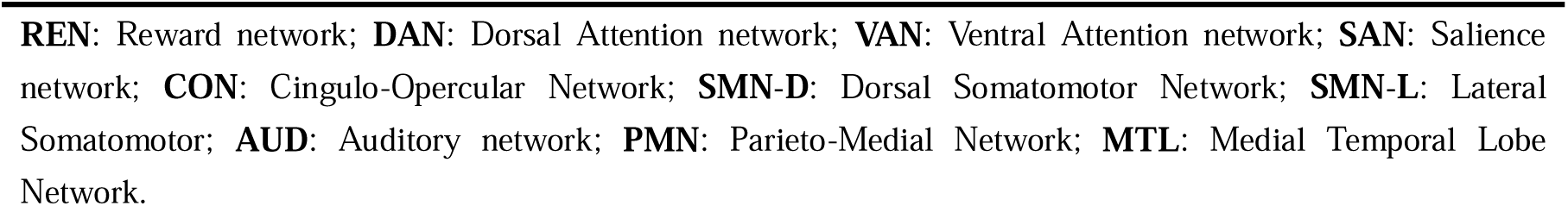
All the seed regions in the network analysis.

| Code | MNI coordinates |  |  | Radius<br>(mm) | Network name | Anatomical |
| --- | --- | --- | --- | --- | --- | --- |
|  | x | y | z |  |  | Labels |
| 1 | -56.16 | -44.76 | -24.23 | 5 | unassigned | cortexL |
| 2 | -24.66 | -97.84 | -12.33 | 5 | unassigned | cortexL |
| 3 | 8.13 | 41.12 | -24.31 | 5 | unassigned | cortexR |
| 4 | 26.68 | -97.3 | -13.49 | 5 | unassigned | cortexR |
| 5 | 48.52 | -2.85 | -38.49 | 5 | unassigned | cortexR |
| 6 | 51.79 | -34.17 | -27.23 | 5 | unassigned | cortexR |
| 7 | 55.18 | -30.8 | -16.93 | 5 | unassigned | cortexR |
| 8 | -57.97 | -25.69 | -14.73 | 5 | unassigned | cortexL |
| 9 | 64.6 | -24.41 | -18.57 | 5 | unassigned | cortexR |
| 10 | -50.06 | -7.09 | -39.24 | 5 | unassigned | cortexL |
| 11 | 33.55 | 38.46 | -12.03 | 5 | unassigned | cortexR |
| 12 | -46.68 | -50.91 | -20.91 | 5 | unassigned | cortexL |
| 13 | -7.12 | -52.22 | 60.71 | 5 | SMN-D | cortexL |
| 14 | 0.05 | -14.53 | 46.74 | 5 | SMN-D | cortexMid |
| 15 | -53.52 | -22.54 | 43.1 | 5 | SMN-D | cortexL |
| 16 | -45.1 | -31.85 | 46.63 | 5 | SMN-D | cortexL |
| 17 | -39.63 | -19.04 | 54.21 | 5 | SMN-D | cortexL |
| 18 | -38.28 | -27.17 | 69.45 | 5 | SMN-D | cortexL |
| 19 | -38.24 | -14.57 | 68.72 | 5 | SMN-D | cortexL |
| 20 | -29.1 | -43 | 60.66 | 5 | SMN-D | cortexL |
| 21 | -22.5 | -30.1 | 72.44 | 5 | SMN-D | cortexL |
| 22 | -20.66 | -31.33 | 60.85 | 5 | SMN-D | cortexL |
| 23 | -16.25 | -45.8 | 73.22 | 5 | SMN-D | cortexL |
| 24 | -13.74 | -17.95 | 39.84 | 5 | SMN-D | cortexL |
| 25 | -12.96 | -17.34 | 74.66 | 5 | SMN-D | cortexL |
| 26 | -6.9 | -20.59 | 65.21 | 5 | SMN-D | cortexL |
| 27 | -6.79 | -33.09 | 72.27 | 5 | SMN-D | cortexL |
| 28 | 2.4 | -27.94 | 60.15 | 5 | SMN-D | cortexR |
| 29 | 3.45 | -17.44 | 58.45 | 5 | SMN-D | cortexR |
| 30 | 9.5 | -1.84 | 44.73 | 5 | SMN-D | cortexR |
| 31 | 9.94 | -45.52 | 72.63 | 5 | SMN-D | cortexR |
| 32 | 10.09 | -17.1 | 74.14 | 5 | SMN-D | cortexR |
| 33 | 13.19 | -32.82 | 74.98 | 5 | SMN-D | cortexR |
| 34 | 20.21 | -28.8 | 59.8 | 5 | SMN-D | cortexR |
| 35 | 22.45 | -42.29 | 68.99 | 5 | SMN-D | cortexR |
| 36 | 28.88 | -16.95 | 70.55 | 5 | SMN-D | cortexR |
| 37 | 28.54 | -39.24 | 59.17 | 5 | SMN-D | cortexR |
| 38 | 37.74 | -17.3 | 45.01 | 5 | SMN-D | cortexR |
| 39 | 42.14 | -20.24 | 54.59 | 5 | SMN-D | cortexR |
| 40 | 44.34 | -7.55 | 56.98 | 5 | SMN-D | cortexR |
| 41 | 47.21 | -29.75 | 48.7 | 5 | SMN-D | cortexR |
| 42 | 50.24 | -20.37 | 41.74 | 5 | SMN-D | cortexR |
| 43 | 19.33 | -7.71 | 63.88 | 5 | SMN-D | cortexR |
| 44 | 54.22 | -27.83 | 33.64 | 5 | SMN-D | cortexR |
| 45 | -52.84 | -10.23 | 24.41 | 5 | SMN-L | cortexL |
| 46 | -49.47 | -11.06 | 34.95 | 5 | SMN-L | cortexL |
| 47 | 36.04 | -9.44 | 13.95 | 5 | SMN-L | cortexR |
| 48 | 51.14 | -5.8 | 32.42 | 5 | SMN-L | cortexR |
| 49 | 65.64 | -7.88 | 24.83 | 5 | SMN-L | cortexR |
| 50 | -51.26 | 8.26 | -2.06 | 5 | CON | cortexL |
| 51 | -44.76 | 0.1 | 8.83 | 5 | CON | cortexL |
| 52 | -34.37 | 3.29 | 4.19 | 5 | CON | cortexL |
| 53 | -16.14 | -4.82 | 70.83 | 5 | CON | cortexL |
| 54 | -10.48 | -2.1 | 42.02 | 5 | CON | cortexL |
| 55 | -5.33 | 17.8 | 34.41 | 5 | CON | cortexL |
| 56 | -2.88 | 2.38 | 53.21 | 5 | CON | cortexL |
| 57 | 6.52 | 7.69 | 50.58 | 5 | CON | cortexR |
| 58 | 13.21 | -1.36 | 69.98 | 5 | CON | cortexR |
| 59 | 35.83 | 10.32 | 1.18 | 5 | CON | cortexR |
| 60 | 36.73 | 0.78 | -3.57 | 5 | CON | cortexR |
| 61 | 49.4 | 8.32 | -1.12 | 5 | CON | cortexR |
| 62 | -30.12 | -27.02 | 12.2 | 5 | CON | cortexL |
| 63 | 42.05 | -0.39 | 47.1 | 5 | CON | cortexR |
| 64 | 35.91 | 21.91 | 2.62 | 5 | CON | cortexR |
| 65 | -39.12 | 50.79 | 17.38 | 5 | CON | cortexL |
| 66 | 31.24 | 32.79 | 26.39 | 5 | CON | cortexR |
| 67 | 31.07 | 55.71 | 14.49 | 5 | CON | cortexR |
| 68 | 36.89 | 32.35 | -2.24 | 5 | CON | cortexR |
| 69 | -52.92 | -21.83 | 22.97 | 5 | AUD | cortexL |
| 70 | 43.45 | -22.93 | 19.85 | 5 | AUD | cortexR |
| 71 | 59.4 | -17.34 | 28.69 | 5 | AUD | cortexR |
| 72 | -60.48 | -25.22 | 13.82 | 5 | AUD | cortexL |
| 73 | -55.22 | -9.42 | 11.73 | 5 | AUD | cortexL |
| 74 | -49.77 | -34.36 | 25.74 | 5 | AUD | cortexL |
| 75 | -49.14 | -26.3 | 5.18 | 5 | AUD | cortexL |
| 76 | -38.43 | -33.34 | 16.98 | 5 | AUD | cortexL |
| 77 | 31.75 | -26.33 | 12.91 | 5 | AUD | cortexR |
| 78 | 55.96 | -5.03 | 13.25 | 5 | AUD | cortexR |
| 79 | 57.88 | -15.62 | 7.49 | 5 | AUD | cortexR |
| 80 | 65.43 | -33.2 | 19.97 | 5 | AUD | cortexR |
| 81 | -68.47 | -22.66 | -15.74 | 5 | DMN | cortexL |
| 82 | -68.3 | -41.41 | -5.14 | 5 | DMN | cortexL |
| 83 | -57.75 | -29.7 | -3.94 | 5 | DMN | cortexL |
| 84 | -55.72 | -12.96 | -10.24 | 5 | DMN | cortexL |
| 85 | -52.89 | 2.55 | -27.06 | 5 | DMN | cortexL |
| 86 | -49.3 | -42.15 | 0.83 | 5 | DMN | cortexL |
| 87 | -45.79 | -60.69 | 20.85 | 5 | DMN | cortexL |
| 88 | -46.17 | 31.26 | -13.03 | 5 | DMN | cortexL |
| 89 | -44.45 | -64.64 | 34.78 | 5 | DMN | cortexL |
| 90 | -43.58 | 11.99 | -34.15 | 5 | DMN | cortexL |
| 91 | -40.5 | -75.27 | 25.8 | 5 | DMN | cortexL |
| 92 | -39.05 | -74.95 | 43.72 | 5 | DMN | cortexL |
| 93 | -35.36 | 19.86 | 50.8 | 5 | DMN | cortexL |
| 94 | -19.78 | 45.07 | 39.48 | 5 | DMN | cortexL |
| 95 | -20.16 | 63.65 | 19.39 | 5 | DMN | cortexL |
| 96 | -17.65 | 63.19 | -9.17 | 5 | DMN | cortexL |
| 97 | -16.4 | 28.52 | 53.05 | 5 | DMN | cortexL |
| 98 | -12.6 | -39.64 | 0.93 | 5 | DMN | cortexL |
| 99 | -11.29 | -56.2 | 15.6 | 5 | DMN | cortexL |
| 100 | -10.09 | 39.09 | 52.29 | 5 | DMN | cortexL |
| 101 | -10.33 | 54.63 | 38.71 | 5 | DMN | cortexL |
| 102 | -11.06 | 44.62 | 7.61 | 5 | DMN | cortexL |
| 103 | -7.55 | 48.08 | 23.18 | 5 | DMN | cortexL |
| 104 | -6.84 | -54.9 | 27.05 | 5 | DMN | cortexL |
| 105 | -7.04 | 50.82 | -1.29 | 5 | DMN | cortexL |
| 106 | -2.94 | -48.79 | 12.87 | 5 | DMN | cortexL |
| 107 | -2.2 | -36.68 | 43.85 | 5 | DMN | cortexL |
| 108 | -3.06 | 44.41 | -9.46 | 5 | DMN | cortexL |
| 109 | -2.06 | 37.85 | 36.34 | 5 | DMN | cortexL |
| 110 | -2.5 | 41.7 | 16.05 | 5 | DMN | cortexL |
| 111 | 5.55 | 66.69 | -3.55 | 5 | DMN | cortexR |
| 112 | 5.91 | -58.82 | 35.45 | 5 | DMN | cortexR |
| 113 | 5.94 | 54.42 | 16.18 | 5 | DMN | cortexR |
| 114 | 6.11 | 63.98 | 21.96 | 5 | DMN | cortexR |
| 115 | 7.94 | -48.37 | 30.57 | 5 | DMN | cortexR |
| 116 | 7.51 | 42.49 | -5.35 | 5 | DMN | cortexR |
| 117 | 8.36 | 47.59 | -15.18 | 5 | DMN | cortexR |
| 118 | 8.8 | 54.23 | 3.45 | 5 | DMN | cortexR |
| 119 | 10.77 | -53.83 | 17.09 | 5 | DMN | cortexR |
| 120 | 12.73 | 54.87 | 38.19 | 5 | DMN | cortexR |
| 121 | 13.08 | 29.99 | 58.65 | 5 | DMN | cortexR |
| 122 | 12.25 | 35.63 | 20.3 | 5 | DMN | cortexR |
| 123 | 15.12 | -63.09 | 25.98 | 5 | DMN | cortexR |
| 124 | 22.11 | 39.21 | 38.9 | 5 | DMN | cortexR |
| 125 | 23.33 | 33.07 | 47.68 | 5 | DMN | cortexR |
| 126 | 43.43 | -72.21 | 28 | 5 | DMN | cortexR |
| 127 | 45.64 | 16.2 | -30.02 | 5 | DMN | cortexR |
| 128 | 46.68 | -50.08 | 28.76 | 5 | DMN | cortexR |
| 129 | 49.26 | 35.47 | -12.2 | 5 | DMN | cortexR |
| 130 | 52.04 | -59.37 | 35.52 | 5 | DMN | cortexR |
| 131 | 52.16 | -2.43 | -16.4 | 5 | DMN | cortexR |
| 132 | 51.9 | 6.81 | -29.61 | 5 | DMN | cortexR |
| 133 | 64.64 | -11.8 | -19.3 | 5 | DMN | cortexR |
| 134 | 64.8 | -30.55 | -8.7 | 5 | DMN | cortexR |
| 135 | -33.93 | -38.06 | -15.6 | 5 | DMN | cortexL |
| 136 | 4.2 | -48.06 | 50.71 | 5 | PMN | cortexR |
| 137 | -6.58 | -71.47 | 41.74 | 5 | PMN | cortexL |
| 138 | -2.47 | -34.8 | 31.07 | 5 | PMN | cortexL |
| 139 | 1.75 | -24.25 | 30.36 | 5 | PMN | cortexR |
| 140 | 11.27 | -66.01 | 42.09 | 5 | PMN | cortexR |
| 141 | -12.08 | -94.56 | -12.8 | 5 | VIS | cortexL |
| 142 | 7.98 | -91.08 | -7.1 | 5 | VIS | cortexR |
| 143 | 17.27 | -91.09 | -13.64 | 5 | VIS | cortexR |
| 144 | -46.54 | -75.95 | -9.95 | 5 | VIS | cortexL |
| 145 | -42.1 | -73.62 | 0.38 | 5 | VIS | cortexL |
| 146 | -40.21 | -88.44 | -6.19 | 5 | VIS | cortexL |
| 147 | -33 | -79.02 | -13.24 | 5 | VIS | cortexL |
| 148 | -28.07 | -79.45 | 19.43 | 5 | VIS | cortexL |
| 149 | -26.39 | -90.23 | 3.12 | 5 | VIS | cortexL |
| 150 | -23.94 | -90.98 | 18.96 | 5 | VIS | cortexL |
| 151 | -17.87 | -68.03 | 4.81 | 5 | VIS | cortexL |
| 152 | -16.21 | -76.97 | 33.82 | 5 | VIS | cortexL |
| 153 | -15.85 | -52.34 | -1.43 | 5 | VIS | cortexL |
| 154 | -15.02 | -72.42 | -7.68 | 5 | VIS | cortexL |
| 155 | -14.22 | -90.66 | 31.4 | 5 | VIS | cortexL |
| 156 | -8.43 | -80.5 | 7.44 | 5 | VIS | cortexL |
| 157 | -2.88 | -81.25 | 21.1 | 5 | VIS | cortexL |
| 158 | 5.59 | -71.65 | 23.52 | 5 | VIS | cortexR |
| 159 | 6.21 | -81.41 | 6.11 | 5 | VIS | cortexR |
| 160 | 8.45 | -71.84 | 10.79 | 5 | VIS | cortexR |
| 161 | 15.27 | -87.09 | 36.89 | 5 | VIS | cortexR |
| 162 | 15.18 | -76.68 | 31 | 5 | VIS | cortexR |
| 163 | 17.53 | -46.86 | -9.88 | 5 | VIS | cortexR |
| 164 | 19.81 | -65.56 | 1.72 | 5 | VIS | cortexR |
| 165 | 19.64 | -85.62 | -2.39 | 5 | VIS | cortexR |
| 166 | 24.41 | -87.21 | 24.01 | 5 | VIS | cortexR |
| 167 | 25.66 | -79.47 | -15.56 | 5 | VIS | cortexR |
| 168 | 26.93 | -59.37 | -9.36 | 5 | VIS | cortexR |
| 169 | 28.68 | -76.62 | 25.42 | 5 | VIS | cortexR |
| 170 | 36.76 | -84.11 | 12.99 | 5 | VIS | cortexR |
| 171 | 36.51 | -81.16 | 1.2 | 5 | VIS | cortexR |
| 172 | 39.98 | -72.49 | 14.36 | 5 | VIS | cortexR |
| 173 | 41.6 | -65.5 | -8.27 | 5 | VIS | cortexR |
| 174 | 42.52 | -78.17 | -11.78 | 5 | VIS | cortexR |
| 175 | -52.6 | -48.83 | 42.5 | 5 | FPN | cortexL |
| 176 | -46.5 | 10.85 | 23.04 | 5 | FPN | cortexL |
| 177 | -43.93 | 1.8 | 45.7 | 5 | FPN | cortexL |
| 178 | -42.23 | 38.21 | 21.35 | 5 | FPN | cortexL |
| 179 | -42.09 | -54.98 | 44.74 | 5 | FPN | cortexL |
| 180 | -42.1 | 24.68 | 29.53 | 5 | FPN | cortexL |
| 181 | -41.06 | 5.81 | 32.72 | 5 | FPN | cortexL |
| 182 | -41.68 | 45.16 | -2.31 | 5 | FPN | cortexL |
| 183 | -34.16 | 54.83 | 4.36 | 5 | FPN | cortexL |
| 184 | -28.4 | -57.93 | 47.78 | 5 | FPN | cortexL |
| 185 | -22.53 | 10.76 | 63.73 | 5 | FPN | cortexL |
| 186 | -2.98 | 26.41 | 44.42 | 5 | FPN | cortexL |
| 187 | 24.07 | 44.61 | -15.35 | 5 | FPN | cortexR |
| 188 | 31.83 | 14.37 | 55.98 | 5 | FPN | cortexR |
| 189 | 33.38 | -53.12 | 44.02 | 5 | FPN | cortexR |
| 190 | 33.6 | 54.22 | -12.95 | 5 | FPN | cortexR |
| 191 | 37.45 | -64.7 | 40.38 | 5 | FPN | cortexR |
| 192 | 38.37 | 43.18 | 15.06 | 5 | FPN | cortexR |
| 193 | 39.87 | 18.39 | 39.72 | 5 | FPN | cortexR |
| 194 | 43.93 | -52.95 | 46.95 | 5 | FPN | cortexR |
| 195 | 43.25 | 49.25 | -2.31 | 5 | FPN | cortexR |
| 196 | 47.01 | 9.93 | 32.66 | 5 | FPN | cortexR |
| 197 | 47.98 | 24.56 | 26.5 | 5 | FPN | cortexR |
| 198 | 49.18 | -42.41 | 45.16 | 5 | FPN | cortexR |
| 199 | 58.31 | -52.79 | -13.61 | 5 | FPN | cortexR |
| 200 | 47.6 | 22.16 | 9.74 | 5 | FPN | cortexR |
| 201 | 55.27 | -44.59 | 36.7 | 5 | FPN | cortexR |
| 202 | -35.44 | 20.03 | 0.07 | 5 | SAN | cortexL |
| 203 | -27.5 | 52.04 | 21.28 | 5 | SAN | cortexL |
| 204 | -10.76 | 25.99 | 24.54 | 5 | SAN | cortexL |
| 205 | -0.94 | 14.86 | 43.99 | 5 | SAN | cortexL |
| 206 | -0.2 | 30.35 | 27.22 | 5 | SAN | cortexL |
| 207 | 5.23 | 23.22 | 37.03 | 5 | SAN | cortexR |
| 208 | 10.26 | 22.06 | 27.48 | 5 | SAN | cortexR |
| 209 | 26.07 | 49.56 | 26.58 | 5 | SAN | cortexR |
| 210 | 33.56 | 16.45 | -7.58 | 5 | SAN | cortexR |
| 211 | 51.28 | -28.52 | -4.3 | 5 | VAN | cortexR |
| 212 | -56.47 | -50.48 | 9.92 | 5 | VAN | cortexL |
| 213 | -55.3 | -39.89 | 13.51 | 5 | VAN | cortexL |
| 214 | 51.52 | -32.52 | 7.55 | 5 | VAN | cortexR |
| 215 | 55.75 | -46.07 | 11.42 | 5 | VAN | cortexR |
| 216 | -49.07 | 25.13 | -0.98 | 5 | VAN | cortexL |
| 217 | -9.88 | 10.95 | 66.61 | 5 | VAN | cortexL |
| 218 | 52.68 | 32.58 | 0.57 | 5 | VAN | cortexR |
| 219 | 53.9 | -42.76 | 21.83 | 5 | VAN | cortexR |
| 220 | -26.6 | -70.72 | 36.86 | 5 | DAN | cortexL |
| 221 | -37.26 | -28.8 | -25.58 | 5 | DAN | cortexL |
| 222 | 45.68 | -46.67 | -16.85 | 5 | DAN | cortexR |
| 223 | 10.51 | -38.54 | 50.02 | 5 | DAN | cortexR |
| 224 | -52.44 | -63.14 | 5.29 | 5 | DAN | cortexL |
| 225 | -42.26 | -60.12 | -8.85 | 5 | DAN | cortexL |
| 226 | -32.56 | -46.42 | 47.2 | 5 | DAN | cortexL |
| 227 | -32.23 | -1.08 | 54.06 | 5 | DAN | cortexL |
| 228 | -16.5 | -58.57 | 64.46 | 5 | DAN | cortexL |
| 229 | 9.61 | -61.5 | 60.88 | 5 | DAN | cortexR |
| 230 | 21.9 | -64.74 | 48.12 | 5 | DAN | cortexR |
| 231 | 25.34 | -58.18 | 60.34 | 5 | DAN | cortexR |
| 232 | 28.56 | -4.62 | 53.99 | 5 | DAN | cortexR |
| 233 | 46.09 | -58.93 | 3.93 | 5 | DAN | cortexR |
| 234 | -31.13 | -9.99 | -36.32 | 5 | MTL | cortexL |
| 235 | 32.85 | -12.41 | -34.41 | 5 | MTL | cortexR |
| 236 | -25.57 | -11.78 | -21.54 | 4 | MTL | hippocampus |
| 237 | 24.73 | -11.25 | -22.68 | 4 | MTL | hippocampus |
| 238 | -25.24 | -38.78 | -2.01 | 4 | DMN | hippocampus |
| 239 | 25.08 | -37.18 | -2.16 | 4 | DMN | hippocampus |
| 240 | -30.63 | 18.71 | -18.98 | 5 | REN | cortexL |
| 241 | -21.14 | 40.87 | -20.48 | 5 | REN | cortexL |
| 242 | 23.96 | 31.94 | -17.78 | 5 | REN | cortexR |
| 243 | 27.06 | 16.22 | -16.93 | 5 | REN | cortexR |
| 244 | -20.3 | -2.27 | -22.21 | 4 | REN | amygdala |
| 245 | 19.51 | -1.85 | -23.11 | 4 | REN | amygdala |
| 246 | 12.66 | 17.32 | -5.06 | 4 | REN | basalGanglia |
| 247 | -12.49 | 17.05 | -4.49 | 4 | REN | basalGanglia |
| 248 | 11.87 | 17.51 | 6.57 | 4 | SAN | basalGanglia |
| 249 | -13.28 | 17.24 | 7.14 | 4 | SAN | basalGanglia |
| 250 | 14.15 | -1.19 | 18.18 | 4 | FPN | basalGanglia |
| 251 | -15.19 | -1.5 | 18.84 | 4 | FPN | basalGanglia |
| 252 | 28.56 | -7.53 | 7.66 | 4 | SMN-D | basalGanglia |
| 253 | -28.02 | -10.23 | 9.07 | 4 | SMN-D | basalGanglia |
| 254 | 28.28 | -6.7 | -5.06 | 4 | SMN-L | basalGanglia |
| 255 | -28.31 | -10.45 | -3.6 | 4 | SMN-L | basalGanglia |
| 256 | 25.26 | 1.75 | -1.26 | 4 | VAN | basalGanglia |
| 257 | -28.26 | -0.97 | -3.09 | 4 | VAN | basalGanglia |
| 258 | 25.46 | 5.05 | 7 | 4 | CON | basalGanglia |
| 259 | -24.83 | 7.67 | 7.95 | 4 | CON | basalGanglia |
| 260 | 18.88 | -4.68 | -3.91 | 4 | CON | basalGanglia |
| 261 | -18.84 | -5.08 | -3.06 | 4 | CON | basalGanglia |
| 262 | 3.42 | -7.79 | 8.23 | 4 | DMN | thalamus |
| 263 | -2.88 | -9.96 | 8.5 | 4 | DMN | thalamus |
| 264 | 18.89 | -28.79 | 0.66 | 4 | VIS | thalamus |
| 265 | -18.84 | -29.18 | 1.51 | 4 | VIS | thalamus |
| 266 | 12.9 | -13.94 | 11.55 | 4 | CON | thalamus |
| 267 | -15.39 | -14.24 | 12.19 | 4 | CON | thalamus |
| 268 | 9.56 | -7.84 | 1.76 | 4 | CON | thalamus |
| 269 | -9.35 | -10.17 | 0.2 | 4 | CON | thalamus |
| 270 | 13.67 | -19.39 | 0.23 | 4 | SMN-L | thalamus |
| 271 | -13.57 | -19.67 | 0.84 | 4 | SMN-L | thalamus |
| 272 | 15.95 | -22.37 | 8.8 | 4 | SMN-D | thalamus |
| 273 | -18.63 | -22.73 | 9.59 | 4 | SMN-D | thalamus |
| 274 | 32 | -49 | -51 | 4 | CON | cerebellum |
| 275 | -13 | -52 | -50 | 4 | DAN | cerebellum |
| 276 | 14 | -48 | -52 | 4 | DAN | cerebellum |
| 277 | -32 | -78 | -38 | 4 | DMN | cerebellum |
| 278 | 32 | -81 | -38 | 4 | DMN | cerebellum |
| 279 | -24 | -76 | -28 | 4 | DMN | cerebellum |
| 280 | 24 | -76 | -28.01 | 4 | DMN | cerebellum |
| 281 | -5.72 | -50.8 | -40.84 | 4 | DMN | cerebellum |
| 282 | 8 | -50 | -40 | 4 | DMN | cerebellum |
| 283 | 0 | -74 | -25 | 4 | VIS | cerebellum |
| 284 | -10 | -78 | -28 | 4 | FPN | cerebellum |
| 285 | 10 | -78 | -28 | 4 | FPN | cerebellum |
| 286 | -34 | -72.01 | -48 | 4 | FPN | cerebellum |
| 287 | 34 | -72 | -48 | 4 | FPN | cerebellum |
| 288 | -30.5 | -66 | -30 | 4 | FPN | cerebellum |
| 289 | 31.68 | -62.83 | -30.4 | 4 | FPN | cerebellum |
| 290 | 40 | -44 | -38 | 4 | FPN | cerebellum |
| 291 | 43.5 | -60 | -30 | 4 | SAN | cerebellum |
| 292 | -43.5 | -60 | -30 | 4 | SAN | cerebellum |
| 293 | -34 | -42 | -44 | 4 | CON | cerebellum |
| 294 | -33 | -51 | -50 | 4 | CON | cerebellum |
| 295 | -6 | -74 | -42 | 4 | SMN-D | cerebellum |
| 296 | 7.5 | -72 | -39 | 4 | SMN-D | cerebellum |
| 297 | -10 | -62 | -18 | 4 | SMN-L | cerebellum |
| 298 | 10 | -62 | -18 | 4 | SMN-L | cerebellum |
| 299 | -12 | -44 | -18 | 4 | SMN-D | cerebellum |
| 300 | 12 | -44 | -18 | 4 | SMN-D | cerebellum |

**Notes: unassigned:** Denotes brain regions lacking a predefined network assignment, which were subsequently omitted from the analysis. **DMN:** Default mode network; **VIS:** Visual network; **FPN:** FrontoParietal network;
**REN:** Reward network; **DAN:** Dorsal Attention network; **VAN:** Ventral Attention network; **SAN:** Salience network; **CON:** Cingulo-Opercular Network; **SMN-D:** Dorsal Somatomotor Network; **SMN-L:** Lateral Somatomotor; **AUD:** Auditory network; **PMN:** Parieto-Medial Network; **MTL:** Medial Temporal Lobe Network.

**Table S2:** Proportion of Zero Connections by Sparsity Threshold.

| Sparsity Threshold | Mean Proportion of Zeros (%) | SD (%) | Decision |
| --- | --- | --- | --- |
| 10% | 20.47 | 6.52 | Excluded |
| 30% | 1.61 | 1.32 | Included |
| 50% | 0.24 | 0.53 | Included |
| 70% | 0.02 | 0.16 | Included |
| 90% | 0 | 0 | Included |

Proportions were calculated from the mean functional connectivity matrices for each participant. A 10% sparsity threshold retains the top 10% of connection weights, and so on.

**Table S3:**
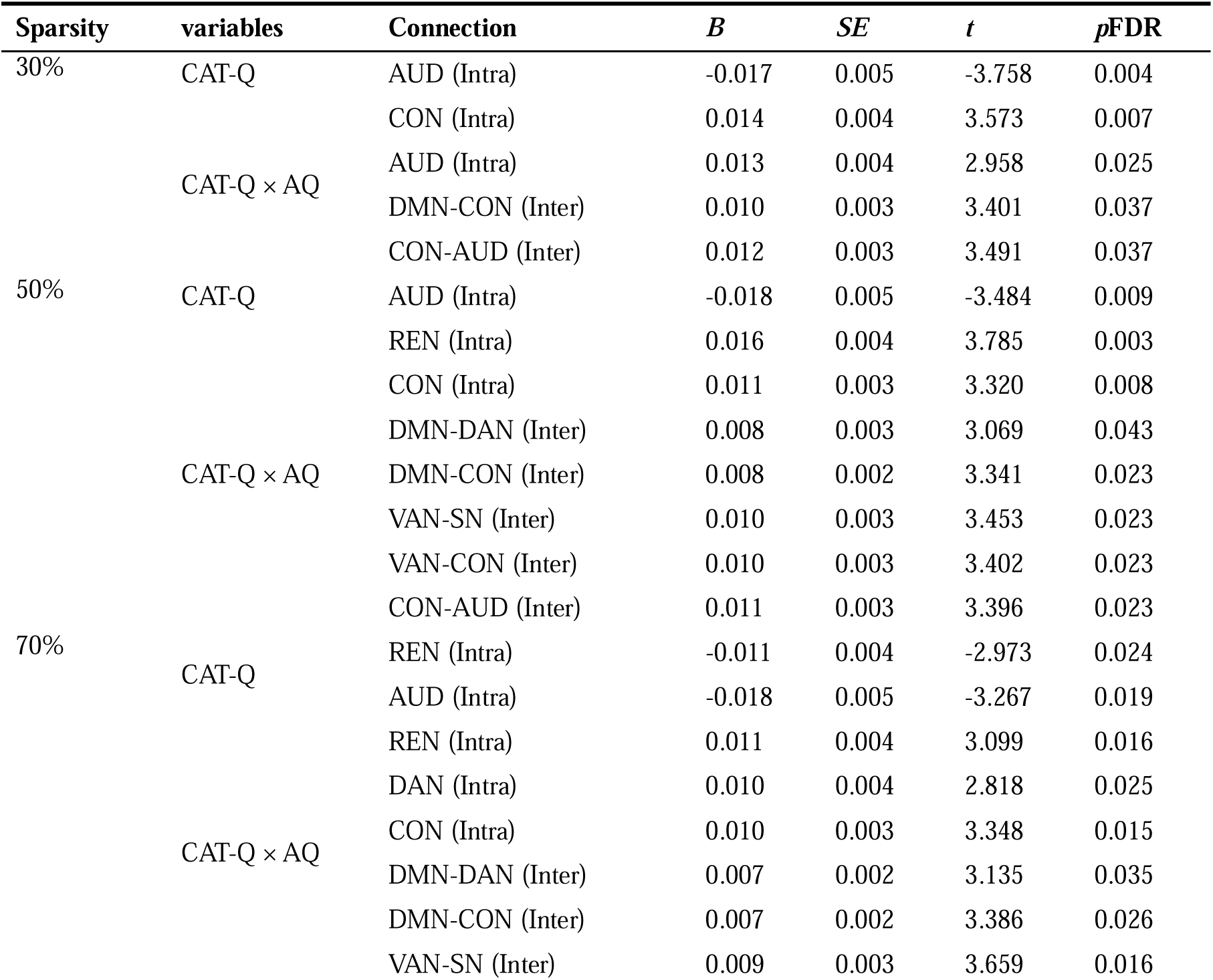
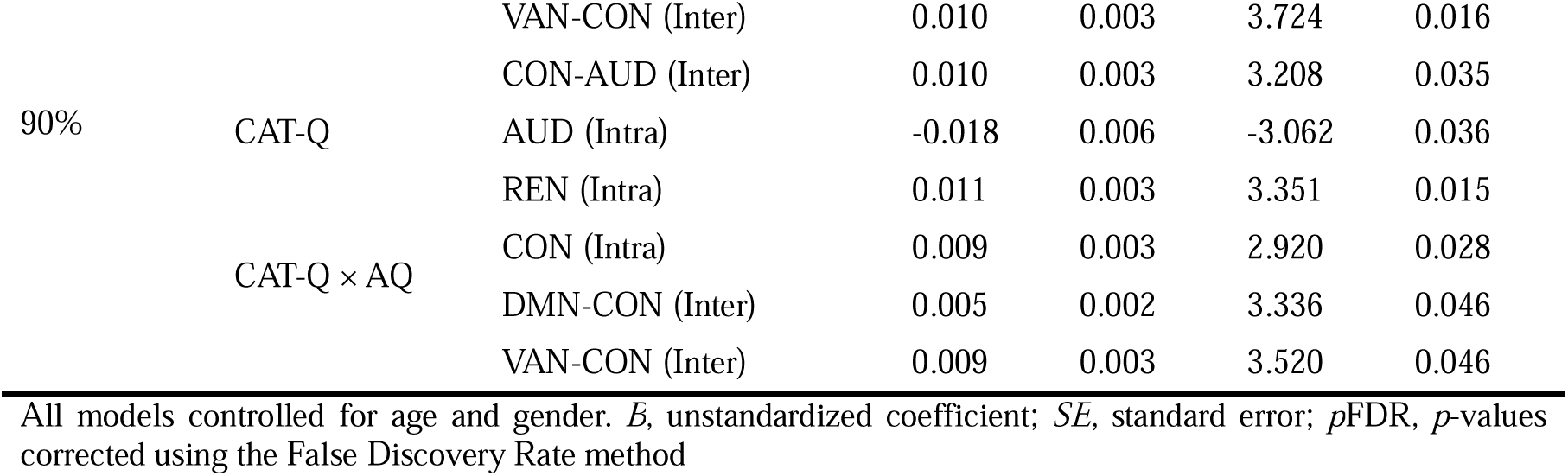
Robustness check for the association between key brain network functional connectivity and scale scores.

**Table S4:** Regression Results for the Moderated Mediation Models (Sparsity 30%)

| Predictors | DMN-CON |  |  |  | TAI |  |  |  | BDI |  |  |  |
| --- | --- | --- | --- | --- | --- | --- | --- | --- | --- | --- | --- | --- |
|  | *B* | *SE* | *t* | *p* | *B* | *SE* | *t* | *p* | *B* | *SE* | *t* | *p* |
| Constant | 0.168 | 1.403 | 0.119 | 0.905 | -0.189 | 1.355 | -0.140 | 0.889 | 0.755 | 1.389 | 0.544 | 0.588 |
| Gender | -0.196 | 0.215 | -0.911 | 0.364 | 0.382 | 0.208 | 1.841 | 0.068 | 0.218 | 0.213 | 1.026 | 0.307 |
| Age | 0.004 | 0.063 | 0.058 | 0.954 | -0.023 | 0.061 | -0.377 | 0.707 | -0.056 | 0.062 | -0.904 | 0.368 |
| CAT-Q | -0.044 | 0.099 | -0.446 | 0.657 | 0.259 | 0.093 | 2.801 | 0.006 | 0.209 | 0.095 | 2.205 | 0.03 |
| AQ | 0.102 | 0.098 | 1.043 | 0.299 |  |  |  |  |  |  |  |  |
| CAT-Q × AQ | 0.323 | 0.095 | 3.401 | 0.001 | | | | | | | | |
| DMN-CON |  |  |  |  | 0.214 | 0.089 | 2.398 | 0.018 | 0.223 | 0.091 | 2.437 | 0.017 |
| R<sup>2</sup> | 0.122 |  |  |  | 0.411 |  |  |  | 0.127 |  |  |  |
| F | 2.884 |  |  | 0.018 | 5.335 |  |  | 0.001 | 3.817 |  |  | 0.006 |
| TAI: Index of moderated mediation |  |  |  |  | BDI: Index of moderated mediation |  |  |  |  |  |  |  |
|  | Index | Boot SE | Boot LLCI | Boot ULCI | Index | Boot SE | Boot LLCI | Boot ULCI |  |  |  |  |
|  | 0.069 | 0.035 | 0.011 | 0.145 | 0.072 | 0.072 | 0.014 | 0.155 |  |  |  |  |

All models controlled for age and gender. *B*, unstandardized coefficient; *SE*, standard error; *pFDR*, *p*-values corrected using the False Discovery Rate method; Index, Index of moderated mediation; LL, lower limit; UL, upper limit; CI, confidence interval.

**Table S5:** Regression Results for the Moderated Mediation Models (Sparsity 50%)

| Predictors | DMN-CON |  |  |  | TAI |  |  |  | BDI |  |  |  |
| --- | --- | --- | --- | --- | --- | --- | --- | --- | --- | --- | --- | --- |
|  | *B* | *SE* | *t* | *p* | *B* | *SE* | *t* | *p* | *B* | *SE* | *t* | *p* |
| Constant | 0.358 | 1.406 | 0.255 | 0.799 | -0.230 | 1.355 | -0.170 | 0.866 | 0.721 | 1.398 | 0.516 | 0.607 |
| Gender | -0.218 | 0.216 | -1.012 | 0.314 | 0.388 | 0.208 | 1.867 | 0.065 | 0.219 | 0.214 | 1.020 | 0.310 |
| Age | -0.004 | 0.063 | -0.063 | 0.950 | -0.021 | 0.061 | -0.351 | 0.726 | -0.055 | 0.063 | -0.872 | 0.385 |
| CAT-Q | -0.056 | 0.099 | -0.569 | 0.799 | 0.263 | 0.093 | 2.836 | 0.005 | 0.212 | 0.096 | 2.223 | 0.028 |
| AQ | 0.094 | 0.098 | 0.955 | 0.342 |  |  |  |  |  |  |  |  |
| CAT-Q × AQ | 0.318 | 0.095 | 3.441 | 0.001 | | | | | | | | |
| DMN-CON |  |  |  |  | 0.216 | 0.089 | 2.418 | 0.017 | 0.196 | 0.092 | 2.131 | 0.035 |
| R<sup>2</sup> | 0.118 |  |  |  | 0.170 |  |  |  | 0.116 |  |  |  |
| F | 2.793 |  |  | 0.021 | 5.364 |  |  | 0.001 | 3.439 |  |  | 0.011 |

| TAI: Index of moderated mediation |  |  |  | BDI: Index of moderated mediation |  |  |  |
| --- | --- | --- | --- | --- | --- | --- | --- |
| Index | Boot SE | Boot LLCI | Boot ULCI | Index | Boot SE | Boot LLCI | Boot ULCI |
| 0.069 | 0.036 | 0.013 | 0.151 | 0.062 | 0.036 | 0.009 | 0.150 |

All models controlled for age and gender. *B*, unstandardized coefficient; *SE*, standard error; *p*FDR, p-values corrected using the False Discovery Rate method; Index, Index of moderated mediation; LL, lower limit; UL, upper limit; CI, confidence interval.

**Table S6:** Regression Results for the Moderated Mediation Models (Sparsity 70%)

| Predictors | DMN-CON |  |  |  | TAI |  |  |  | BDI |  |  |  |
| --- | --- | --- | --- | --- | --- | --- | --- | --- | --- | --- | --- | --- |
|  | *B* | *SE* | *t* | *p* | *B* | *SE* | *t* | *p* | *B* | *SE* | *t* | *p* |
| Constant | 0.557 | 1.407 | 0.396 | 0.693 | -0.248 | 1.366 | -0.182 | 0.856 | 0.709 | 1.410 | 0.503 | 0.616 |
| Gender | -0.261 | 0.216 | -1.206 | 0.231 | 0.391 | 0.210 | 1.861 | 0.066 | 0.219 | 0.217 | 1.010 | 0.315 |
| Age | -0.010 | 0.064 | -0.164 | 0.870 | -0.021 | 0.061 | -0.337 | 0.736 | -0.054 | 0.063 | -0.855 | 0.394 |
| CAT-Q | -0.031 | 0.099 | -0.311 | 0.757 | 0.260 | 0.093 | 2.782 | 0.006 | 0.210 | 0.096 | 2.179 | 0.032 |
| AQ | 0.047 | 0.098 | 0.478 | 0.634 |  |  |  |  |  |  |  |  |
| CAT-Q × AQ | 0.323 | 0.095 | 3.386 | 0.001 |  |  |  |  |  |  |  |  |
| DMN-CON |  |  |  |  | 0.180 | 0.090 | 1.996 | 0.048 | 0.154 | 0.093 | 1.657 | 0.101 |
| R<sup>2</sup> | 0.116 |  |  |  | 0.155 |  |  |  | 0.101 |  |  |  |
| F | 2.742 |  |  | 0.023 | 4.832 |  |  | 0.001 | 2.952 |  |  | 0.023 |

| TAI: Index of moderated mediation |  |  |  | BDI: Index of moderated mediation |  |  |  |
| --- | --- | --- | --- | --- | --- | --- | --- |
| Index | Boot SE | Boot LLCI | Boot ULCI | Index | Boot SE | Boot LLCI | Boot ULCI |
| 0.058 | 0.035 | 0.005 | 0.137 | 0.050 | 0.032 | 0.001 | 0.126 |

All models controlled for age and gender. *B*, unstandardized coefficient; *SE*, standard error; *p*FDR, p-values corrected using the False Discovery Rate method; Index, Index of moderated mediation; LL, lower limit; UL, upper limit; CI, confidence interval.

**Table S7:** Regression Results for the Moderated Mediation Models (Sparsity 90%)

| Predictors | DMN-CON |  |  |  | TAI |  |  |  | BDI |  |  |  |
| --- | --- | --- | --- | --- | --- | --- | --- | --- | --- | --- | --- | --- |
|  | *B* | *SE* | *t* | *p* | *B* | *SE* | *t* | *p* | *B* | *SE* | *t* | *p* |
| Constant | 1.508 | 1.399 | 1.078 | 0.284 | -0.378 | 1.379 | -0.274 | 0.785 | 0.601 | 1.421 | 0.423 | 0.673 |
| Gender | -0.403 | 0.215 | -1.875 | 0.064 | 0.407 | 0.213 | 1.910 | 0.059 | 0.232 | 0.220 | 1.057 | 0.293 |
| Age | -0.046 | 0.063 | -0.728 | 0.468 | -0.016 | 0.062 | -0.252 | 0.801 | -0.050 | 0.064 | -0.783 | 0.436 |
| CAT-Q | -0.024 | 0.098 | -0.246 | 0.806 | 0.260 | 0.094 | 2.772 | 0.007 | 0.210 | 0.097 | 2.175 | 0.032 |
| AQ | 0.014 | 0.098 | 0.145 | 0.885 |  |  |  |  |  |  |  |  |
| CAT-Q × AQ | 0.316 | 0.095 | 3.336 | 0.001 |  |  |  |  |  |  |  |  |
| DMN-CON |  |  |  |  | 0.153 | 0.092 | 1.670 | 0.098 | 0.129 | 0.094 | 1.366 | 0.175 |
| R<sup>2</sup> | 0.126 |  |  |  | 0.146 |  |  |  | 0.094 |  |  |  |
| F | 3.011 |  |  | 0.014 | 4.491 |  |  | 0.002 | 2.714 |  |  | 0.034 |

| TAI: Index of moderated mediation |  |  |  | BDI: Index of moderated mediation |  |  |  |
| --- | --- | --- | --- | --- | --- | --- | --- |
| Index | Boot SE | Boot LLCI | Boot ULCI | Index | Boot SE | Boot LLCI | Boot ULCI |
| 0.048 | 0.033 | -0.001 | 0.123 | 0.041 | 0.029 | -0.006 | 0.107 |

All models controlled for age and gender. *B*, unstandardized coefficient; *SE*, standard error; *p*FDR, p-values corrected using the False Discovery Rate method; Index, Index of moderated mediation; LL, lower limit; UL, upper limit; CI, confidence interval.

## Notes

### Competing Interest Statement

The authors have declared no competing interest.

